# Spectral harmonicity distinguishes two types of ictal phase-amplitude cross frequency couplings in patients candidate to epilepsy surgery

**DOI:** 10.1101/2020.03.13.991299

**Authors:** Damián Dellavale, Eugenio Urdapilleta, Nuria Cámpora, Osvaldo Matías Velarde, Silvia Kochen, Germán Mato

## Abstract

**Objective:** Spectral harmonicity of the ictal activity was analyzed regarding two clinically relevant aspects, (1) as a confounding factor producing ‘spurious’ phase-amplitude couplings (PAC) which may lead to wrong conclusions about the underlying ictal mechanisms, and (2) its role in how good PAC is in correspondence to the seizure onset zone (SOZ) classification performed by the epileptologists.

**Methods:** PAC patterns observed in intracerebral electroencephalography (iEEG) recordings were retrospectively studied during seizures of seven patients with pharmacoresistant focal epilepsy. The time locked index (TLI) measure was introduced to quantify the degree of harmonicity between frequency bands associated to the emergence of PAC during epileptic seizures.

**Results:** (1) Harmonic and non harmonic PAC patterns coexist during the seizure dynamics in iEEG recordings with macroelectrodes. (2) Harmonic PAC patterns are an emergent property of the periodic non sinusoidal waveform constituting the epileptiform activity. (3) The TLI metric allows to distinguish the non harmonic PAC pattern, which has been previously associated with the ictal core through the paroxysmal depolarizing shifts mechanism of seizure propagation.

**Conclusions:** Our results suggest that the spectral harmonicity of the ictal activity plays a relevant role in the visual analysis of the iEEG recordings performed by the epileptologists to define the SOZ, and that it should be considered for the proper interpretation of ictal mechanisms.

**Significance:** The proposed harmonicity analysis can be used to improve the delineation of the SOZ by reliably identifying non harmonic PAC patterns emerging from fully recruited cortical and subcortical areas.

**Highlights:**

- Harmonic and non harmonic phase-amplitude couplings coexist in epileptiform local field potentials.
- The time locked index quantitatively differentiates harmonic and non harmonic ictal PAC patterns.
- Spectral harmonicity plays a relevant role in the interpretation of underlying ictal mechanisms.

## 1. INTRODUCTION

At a macroscopic scale, the electrical activity of the human brain is generally characterized by the presence of multiple oscillatory dynamics originated by spatially distributed ensembles of neurons that interact with each other. Besides, the brain states are commonly observed through neural recordings that project the local network activity into one dimensional time series. In these recordings, the interactions between the underlying oscillatory dynamics is presumably translated to cross frequency couplings (CFC) Canolty et al. (2006); Hyafil et al. (2015) whose intensity varies in a state dependent manner. Thus, the CFC patterns observed at the signal level are hypothesized to be informative on both the underlying physiological neuronal processing Canolty and Knight (2010); Voytek et al. (2010); Nagasawa et al. (2012); Nakatani et al. (2014); Vaz et al. (2017); Bergmann and Born (2018) and the aberrant activity associated to pathological brain states Nariai et al. (2011); Weiss et al. (2013); Ibrahim et al. (2014); Weiss et al. (2015); Edakawa et al. (2016); Zhang et al. (2017); Motoi et al. (2018). From the signal processing point of view, a CFC pattern emerges when certain characteristics (e.g. amplitude, phase) of a frequency band interact with others in a different band, either in the same signal or in another related one. This has motivated the development of specialized signal processing algorithms to robustly detect and quantify CFC phenomena from noisy neural recordings Canolty et al. (2006); Tass et al. (1998); Penny et al. (2008); Cohen (2008); Tort et al. (2010); Cole and Voytek (2017). In spite of these advances, the interpretation of the CFC patterns observed at the signal level remains challenging since it implies to tackle an inverse problem to infer the underlying multidimensional neural dynamics based on the analysis of spatially sparse, one dimensional recordings Velarde et al. (2019). The basic mechanistic interpretation behind CFC is that independent oscillations interact hierarchically Hyafil et al. (2015); Lozano-Soldevilla et al. (2016). For instance, for CFC in the form of phase-amplitude coupling (PAC) Jensen and Colgin (2007), one high frequency (HF) oscillation presumably a manifestation of neural processing in local networks, is amplitude modulated by the phase of a low frequency (LF) oscillation, whose dynamics is governed by the synchronization of large, spatially extended, neuronal ensembles Lozano-Soldevilla et al. (2016). Nevertheless, some fundamental limitations have been identified in connection with this view. The first concern is related to whether the CFC patterns obtained from a given recording reflect a true mechanistic interaction between two independent neural oscillators, or it might be a more trivial consequence of spectral correlations due to some repetitive non sinusoidal waveform constituting the recorded neural activity Cole and Voytek (2017); Lozano-Soldevilla et al. (2016). A second concern refers to whether the CFC pattern reflects a mechanistic causal coupling between the features involved (phase, amplitude, frequency) or it is just an indirect correlation (i.e. an epiphenomenon). Thus, the proper characterization of these nonlinear (cross frequency) interactions observed at the signal level is crucial to identify the underlying mechanisms and its functional significance, which are unsolved issues still under debate in the context of neural processes. On the other hand, ictal PAC patterns involving different frequency bands have been proposed as biomarkers to identify the seizure onset zone (SOZ) and to characterize the seizure dynamics Nariai et al. (2011); Weiss et al. (2013); Ibrahim et al. (2014); Weiss et al. (2015); Edakawa et al. (2016); Zhang et al. (2017); Motoi et al. (2018). Importantly, it has been hypothesized that ictal PAC, compared to interictal PAC, could provide more direct information to delineate the brain regions responsible for generation of habitual seizures Motoi et al. (2018). However, to our knowledge the relationship between spectral harmonicity and ictal PAC, and the possible mechanisms mediating this connection have not been addressed in literature. As a consequence, the question of whether the observed ictal PAC patterns are an epiphenomenon of the (quasi)periodic non sinusoidal waveform shapes constituting the seizure activity or they are related to the interaction of independent neural oscillators remains open.

In this work we perform a quantitative analysis of the ictal PAC patterns in connection with the harmonic spectral content of the seizure activity recorded with clinical macroelectrodes. To this purpose, a novel measure termed time locked index (TLI) was introduced to efficiently quantify the degree of harmonicity between frequency bands associated to the emergence of PAC patterns. Based on this measure, together with others characterizing raw PAC, we provide substantial evidence about the spectral harmonics content of repetitive non sinusoidal waveform constituting the seizure activity as a mechanism originating PAC, referred as ‘harmonic PAC’. Implications of the coexistence of harmonic and non harmonic PAC patterns during epileptic seizures regarding underlying neural mechanisms and improved therapy are also discussed.

## 2. METHODS

### 2.1. Patients

The seven patients with pharmacoresistant focal epilepsy included in this retrospective study were consecutively selected from the candidates for surgical treatment between 2012 and 2015 at the Epilepsy Center - “Ramos Mejía” Hospital and El Cruce “N. Kirchner” Hospital - Universidad Nacional Arturo Jauretche, Argentina. The patients underwent chronic invasive EEG procedure guided by clinical criteria to help identify the epileptogenic zone (EZ) for subsequent resection. The characteristics of the patients are provided in Table 1 and the schematic view of the SOZ, as identified by the epileptologists (NC and SK), is shown in Figure 1.

**Figure 1:**
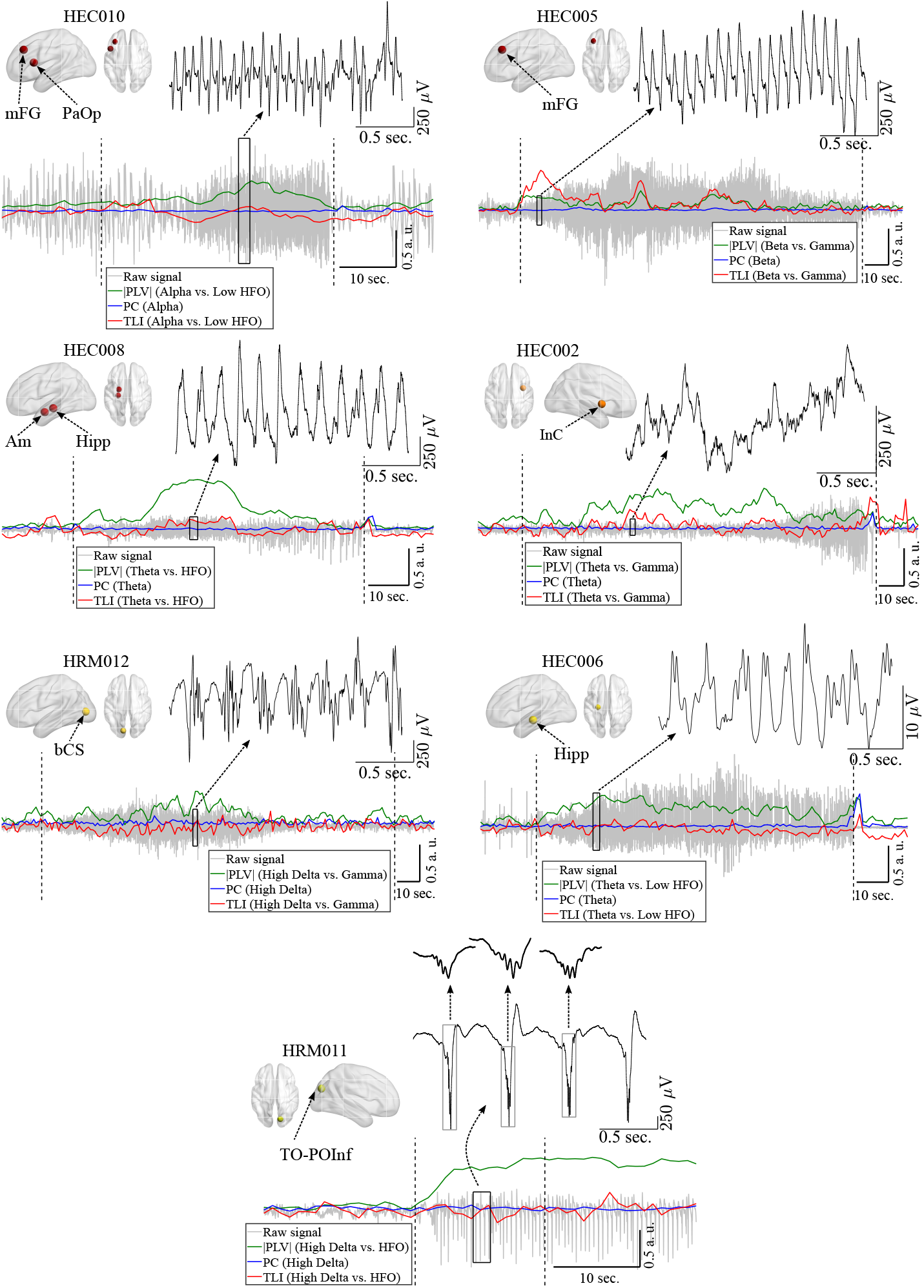
Schematic view of the seizure onset zone (SOZ) localization, together with the bipolar intracerebral electroencephalography (iEEG) recordings (gray line) obtained from the SOZ of each patient (1 bipolar channel per patient). The time marks for the onset and termination of each seizure are indicated by the vertical dashed black lines. The insets (solid black line) show 2 sec. epochs to highlight the oscillatory waveform shapes and time scales during ictal events. The phase locking value (PLV) quantifying phase-amplitude coupling (green line), time locked index (TLI) quantifying harmonicity (red line) and phase clustering (PC) (blue line) time series were constructed using a sliding epoch of 5 sec. in length with 80 % overlap (see Section 2.7). In all the cases, the offset of the TLI time series was removed by substracting the mean value of the pre-ictal TLI time series computed from a time interval of approximately 20 sec. in length located just before the seizure onset. In the case HRM011 (bottom graph), note the non sinusoidal pulse waveforms of the slow rhythm and the fast oscillations at the bottom of negative peaks. Symbols and abbreviations: HFO, high frequency oscillations; a. u., arbitrary units; mFG, middle Frontal Gyrus; PaOp, Pars Opercularis; Am, Amygdala; Hipp, Hippocampus; InC, Insular Cortex; bCS, below Calcarine Sulcus; TO-POInf, Temporo Occipital - Parieto Occipital Inferior.

**Table 1:** Clinical characteristics of the patients.

| Patient<br>Age(ASO) [y]<br>Sex | MRI<br>PET hyp<br>Pathol anat | iEEG location;<br>Elec type(No*) | iEEG SOZ | Region<br>Surgery | Outcome<br>Engel class<br>(Follow-up) | Sz number**<br>Total(Ind) | Sz duration<br>Mean(Std) [s] |
| --- | --- | --- | --- | --- | --- | --- | --- |
| HEC002<br>22(17)<br>Female | Negative<br>r InC<br>N/A | r F T; DM(44) | r InC | r InC<br>cortectomy | IIc<br>(3 years) | 7(0) | 104(50) |
| HEC005<br>19(5)<br>Male | l F<br>N/A<br>CavAng | l F T; DM(45) | l mFG | l mFG<br>lobectomy | I<br>(2 years) | 7(0) | 104(40) |
| HEC006<br>19(18)<br>Male | Negative<br>Diffuse<br>N/A | l/r T; DM(54) | l H/B Hipp | l<br>AH | II<br>(5 years) | 6(2) | 99(29) |
| HEC008<br>36(24)<br>Male | Negative<br>r T / l P O<br>ULHC | l/r M T; DM(54) | l Am Hipp | l<br>AH | IIc<br>(2 years,<br>8 months) | 4(0) | 85(14) |
| HEC010<br>33(7)<br>Male | l F<br>Negative<br>N/A | l F T; DM(39) | l mFG/PaOp | N/A | N/A | 10(0) | 47(9) |
| HRM011<br>20(6)<br>Male | Negative<br>N/A<br>CCD | r T P; DM(14)<br>r T P; Str(8)<br>r F T P O; Grd(74) | r TO POInf | r O<br>cortectomy | IB<br>(4 years) | 9(0) | 50(86) |
| HRM012<br>29(8)<br>Female | Negative<br>N/A<br>CCD | l F T P O; DM(70) | l O bCS | l O<br>cortectomy | IB<br>(4 years) | 3(1) | 180(86) |

Abbreviations: ASO, Age of Seizure Onset; Sz, seizure; y, years; s, seconds; l, left; r, right; M, Medial; H, Head; B, Body; MRI, Magnetic Resonance Imaging; PET hyp, Positron Emission Tomography hypometabolism; Pathol anat, Pathological anatomy; iEEG, intracerebral Electroencephalography; iEEG SOZ, Seizure Onset Zone identified from the intracerebral Electroencephalography recordings; Elec type, Electrode type; Std, Standard deviation; F, Frontal; T, Temporal; P, Parietal; O, Occipital; TO, Temporo-Occipital; POInf, Parieto-Occipital Inferior; InC, Insular Cortex; mFG, middle Frontal Gyrus; Am, Amygdala; Hipp, Hippocampus; PaOp, Pars Opercularis; bCS, below Calcarine Sulcus; CavAng, Cavernous Angioma; ULHC, Undifferentiated Layers in Hippocampal Cortex; CCD, Cytoarchitectural Cortical Dysplasia; AH, Amygdalohippocampectomy; DM, Deep Macroelectrode; Str, Strip; Grd: Grid; N/A, Not Applicable; Total, Total number of seizures analyzed (Spontaneous and Induced); Ind, Number of seizures induced by electrostimulation. \* No, Number of implanted unipolar electrode contacts; \*\* Number of seizures analyzed in this study;

### 2.2. Localization of electrode sites and imaging for three-dimensional reconstruction

Electrode placement considered not only the final target but also the precise trajectory. Surgical planning of electrode placement in specific anatomical structures was thus dependent on the pre-established hypotheses of probable zone of seizure onset and propagation. Pre-implantation hypotheses were determined by two epileptologists (NC and SK) based on seizure semiology, interictal and ictal findings of scalp video-EEG, magnetic resonance imaging data (MRI 3.0 T), positron emission tomography (PET) when was available, together with neuropsychological and psychiatric evaluation. Therefore, electrode positions were not standardized. The exact localization of each electrode was determined using a free and open source medical image computing platform for biomedical research: 3DSlicer and Freesurfer Princich et al. (2013). 3DSlicer enables the fusion of pre-implant MRI and post-implant Computed Tomography (CT) and Freesurfer produces an anatomical parcellation of the cortex. Depth electrodes (Ad-Tech Medical Instrument Corp., Racine, WI, U.S.A.) had: (a) 8 or 10 platinum contacts with 5 or 10 mm inter-contact center to center distance, contact length of 2.4 mm and 1.1 mm diameter, or (b) 9 platinum contacts, 3 mm distance between the first and the second contact and 5 mm inter-electrode distance from the second to the last. Contact length was 1.57 mm and the electrode diameter was 1.28 mm. Electrodes were identified by a letter of the alphabet with no standard labeling for each location. Contacts within a depth electrode were identified with numbers beginning from the deepest located in the tip (contact number 1) to the base of the electrode. Subdural strips and grid electrodes, together with depth electrodes, were used in one of the seven patients (HRM011, see Table 1).

### 2.3. Intracerebral recordings

The intracerebral electroencephalography (iEEG) signals were acquired using an Elite system (Blackrock Neuromed, Utah, U.S.A.), sampled at 2 kHz. The software Cervello 1.04.200 (Micromed, Italy) was used for visualization and analysis of the recordings by the epileptologists. The onset, spread and termination of the seizures were independently identified by two epileptologists (NC and SK). Ictal onset was defined as initial iEEG changes, characterized by sustained rhythmic discharges or repetitive spike-wave discharges that cannot be explained by state changes, and that resulted in habitual seizure symptoms similar to those reported in previous studies. The SOZ in each patient was defined by the epileptologists (NC and SK) as the contacts where the earlier ictal iEEG changes were seen.

### 2.4. Surgical decision making and seizure outcome

The surgical resection area was determined on an individualized basis by the clinical factors, semiology, visual assessment of scalp EEG, MRI data, neuropsychology, psychiatric evaluation and visual assessment of iEEG recordings. The ablative brain surgery was intended to completely remove the SOZ, i.e. iEEG sites with the seizure onset, preserving the eloquent areas and their associated vascular structures along sulcal boundaries. One (HEC010) of the seven patients included in Table 1 did not undergo resective epilepsy surgery after the iEEG study. The rest six patients underwent epilepsy surgery and showed Engel class I-II outcomes following resection.

### 2.5. Signal processing for the retrospective post-hoc analysis

A total of 46 seizures from seven patients were analyzed. Visual inspection of the macroelectrode recordings was made to identify noisy contacts. The local field potentials (LFP) with excessive artifactual content were excluded from this analysis. Overall, across the seven patients, 402 iEEG macroelectrode contacts (i.e. LFP time series) were converted to a bipolar referencing montage for subsequent analysis. Bipolar iEEG channels were obtained as the difference between LFP recorded from spatially adjacent contacts pertaining to the same depth electrode array. In the case of strip and grid electrode matrices each contact was referenced to a nearby contact, moving through the subdural matrix elements along one single dimension (columns or rows). To quantify the PAC dynamics during the seizure activity, non-parametric methods were used: the phase locking value (PLV) and the modulation index based on the Kullback-Leibler distance (KLMI) Tort et al. (2010). Figure 2 shows, for two patients (HEC005 and HEC002), the epochs of ictal LFP (*x*(*t*)) together with the band-pass filtered signals (*x_LF_* (*t*), *x_HF_* (*t*)), the phase of the low frequency signal (*φ_LF_* (*t*)) and, the amplitude envelope of the high frequency signal (*a_HF_* (*t*)), as well as its phase evolution (*φ_aHF_* (*t*)) from which the PLV and KLMI metrics can be computed as follows Penny et al. (2008); Tort et al. (2010),

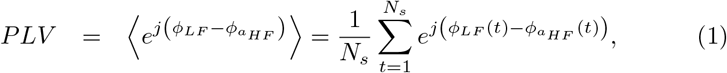

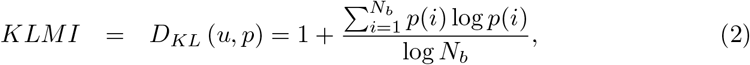

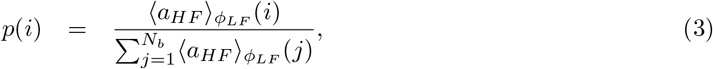

where *N_s_* is the number of samples of the time series, *p*(*i*) denotes the mean amplitude of the high frequency signal at the low frequency phase bin *i* (〈*a_HF_*〉*_φLF_* (*i*)) normalized by the sum over the bins (see histograms in Figure 2), *N_b_* is the number of bins for the phase histogram and *D_KL_* represents the Kullback-Leibler distance between *p* and the uniform distribution *u*. The amplitude envelope (*a*(*t*)) and phase (*φ*(*t*)) time series, required to assess CFC, were computed using the Filter-Hilbert method (see Chapter 14 in Cohen (2014)). In brief, the raw LFP time series *x*(*t*) was band-pass filtered around the frequency band of interest, then, the analytic signal corresponding to the filtered time series was constructed via the Hilbert transform. The *a*(*t*) and *φ*(*t*) for that particular frequency band were obtained by computing the absolute value and argument of the analytic signal, respectively. A detailed description of the method including the computation of the analytic signal in the frequency domain is given in Dellavale and Rosselló (2019). The band-pass filters (BPF) were implemented in the frequency domain by multiplying the Fourier transform of the input signal by a Hann window and then, applying the inverse Fourier transform to get the band-pass filtered signal back in the time domain (i.e. circular convolution in the discrete time domain). Note that this filtering approach was used to effectively isolate the desired frequency bands (i.e. null-to-null bandwidth), which is not guaranteed when other linear filters are used (e.g. low order IIR filters) He et al. (2010). We verified that our BPF implementation do not produce neither phase distortions nor significant oscillations in the output signal capable to generate spurious CFC Widmann and Schröger (2012), showing a performance comparable to that of the FIR filters implemented in the EEGLAB (*eegfilt* function, data not shown) Delorme and Makeig (2004). One of the main confounds when assessing PAC is related to the nonuniform distribution of phase angles of the modulating component *x_LF_* (*t*), which can produce spurious PAC levels van Driel et al. (2015). To detect the occurrence of this confound we computed the phase clustering (PC) as shown in Eq. 4 (see Chapter 30, p. 414 of Cohen (2014)).

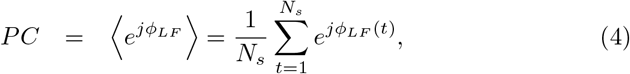

**Figure 2:**
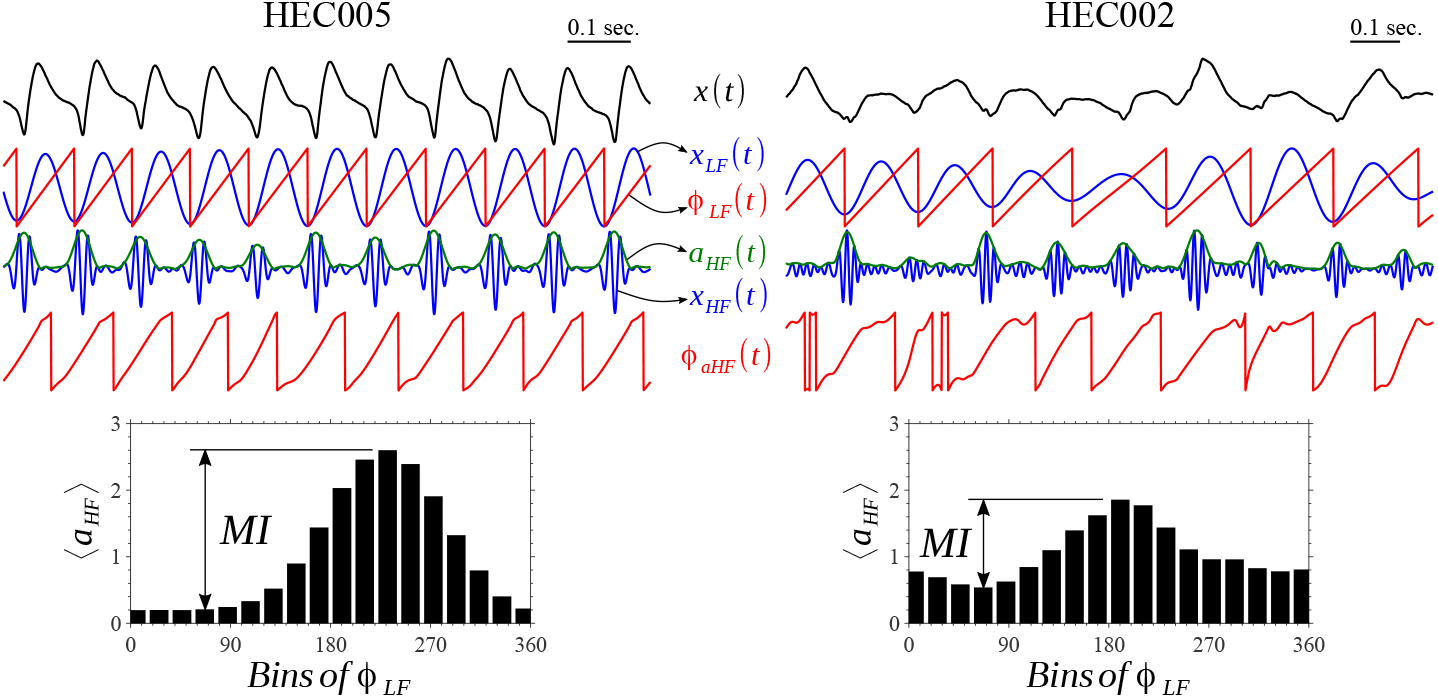
Ictal recordings (bipolar channels) obtained from the seizure onset zone (SOZ) and derived time series involved in the phase locking value (PLV) and modulation index based on the Kullback-Leibler distance (KLMI) algorithms for quantification of phase-amplitude coupling (PAC). The low frequency (LF) band-pass filtered signal (*x_LF_* (*t*)) correspond to the Alpha and Theta bands for patients HEC005 and HEC002, respectively. The high frequency (HF) band-pass filtered signal (*x_HF_* (*t*)) correspond to the Gamma band for both patients. The phase-amplitude histograms at the bottom show that the mean amplitude of the fast oscillations depart from the uniform distribution revealing the existence of a preferred LF phase. Using a time series of 7 sec. in length which included the epoch shown in these plots we obtained PLV = 0.89 (KLMI = 0.13) and PLV = 0.41 (KLMI = 0.024) for patients HEC005 and HEC002, respectively. Symbols and abbreviations: *MI*, modulation index; *x*(*t*), ictal LFP; *φ_LF_* (*t*), phase time series of the *x_LF_* (*t*) signal; *a_HF_* (*t*), amplitude envelope of the *x_HF_* (*t*) signal; *φa_HF_* (*t*), phase time series of the *a_HF_* (*t*) signal.

When *x_LF_* (*t*) has a periodic sinusoidal-like waveform shape, we obtain a rather uniform phase angle distribution *φ_LF_* (*t*) resulting in PC *≈* 0 for a sufficiently large number of samples *N_s_*. On the other hand, if the time series *x_LF_* (*t*) is highly non sinusoidal, a skewed distribution of phase angles is obtained producing PC 1. Worthy to note, the spurious PAC associated to high PC values can be mitigated by using narrow enough BPF to obtain the modulating low frequency oscillations *x_LF_* (*t*) (see the time series *φ_LF_* (*t*) in Figure 2), or by the method described in van Driel et al. (2015). In contrast, to effectively assess PAC, the BPF aimed to obtain the modulated high frequency oscillations *x_HF_* (*t*) must satisfy the restriction related to the minimum bandwidth (*Bw_HF_*) determined by the low frequency band: *Bw_HF_* ≳ 2*f_LF_*, where *f_LF_* is the center frequency of the BPF for *x_LF_* (*t*) Berman et al. (2012).

Time locked plots (TLPs) were used as a complementary tool to analyze the link between the waveform shapes and PAC patterns observed during the seizure dynamics. The TLPs were computed as described in Refs. Canolty et al. (2006); Tort et al. (2008). Fourier spectrograms were computed using Thomson multitaper and modified periodogram methods with a Hann window in the time domain Cohen (2014).

### 2.6. Time locked index

A specialized tool was developed to characterize the ictal PAC patterns observed across the patients Velarde et al. (2019). Specifically, the time locked index (TLI) was implemented to efficiently quantify the presence of spectral harmonics associated to the emergence of PAC in noisy signals and, in particular, in ictal neural recordings. The quantitative characterization of the harmonicity of the ictal activity is important given that coupled oscillatory dynamics with independent frequencies or non sinusoidal brain activity can both elicit a similar signature in the Fourier spectrum. In particular, the traditional algorithms aimed to assess PAC (e.g. PLV, KLMI) are confounded by harmonically related spectral components associated to non sinusoidal brain activity, reporting significant PAC levels in absence of independent frequency bands Vaz et al. (2017); Velarde et al. (2019); Lozano-Soldevilla et al. (2016). In the TLI algorithm, time-locked averages are implemented in the time domain to exploit the phase synchronization between harmonically related spectral components constituting the non sinusoidal oscillatory dynamics. The following steps describe the procedure to compute TLI (see Figure 3),

1. The input signal *x* is band-pass filtered at the low (LF) and high (HF) frequency bands under analysis, producing the time series *x_LF_* and *x_HF_*, respectively. Z-score normalization is applied on the time series *x_LF_* and *x_HF_* to ensure the TLI metric is independent of the signals amplitude.
2. The time instants corresponding to the maximum amplitude (or any other particular phase) of both time series, *x_LF_* and *x_HF_*, are identified in each period of the low frequency band (*T_LF_*). These time values for the slow and fast oscillation peaks are recorded in the time vectors *t_LF_* (red down-pointing triangles in Figure 3) and *t_HF_* (green up-pointing triangles in Figure 3), respectively.
3. Epochs 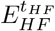 with a length equal to one period of the low frequency band (*T_LF_*) centered at the fast oscillation peaks (*t_HF_*) are extracted form the time series *x_HF_*. Averaging over these epochs is computed to produce a mean epoch 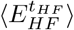. Note that the latter is a time-locked averaging due to the fact that every single epoch 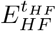 is centered at the corresponding time instant *t_HF_*.
4. Epochs 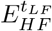 with a length equal to one period of the low frequency band (*T_LF_*) centered at slow oscillation peaks (*t_LF_*) are extracted form the time series *x_HF_*. Averaging over these epochs is computed to produce a mean epoch 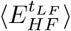. Note that the latter is also a time-locked averaging, now with epochs centered at the corresponding time instants *t_LF_*.
5. Finally, the TLI is computed as follows,

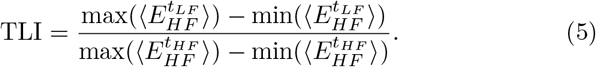

**Figure 3:**
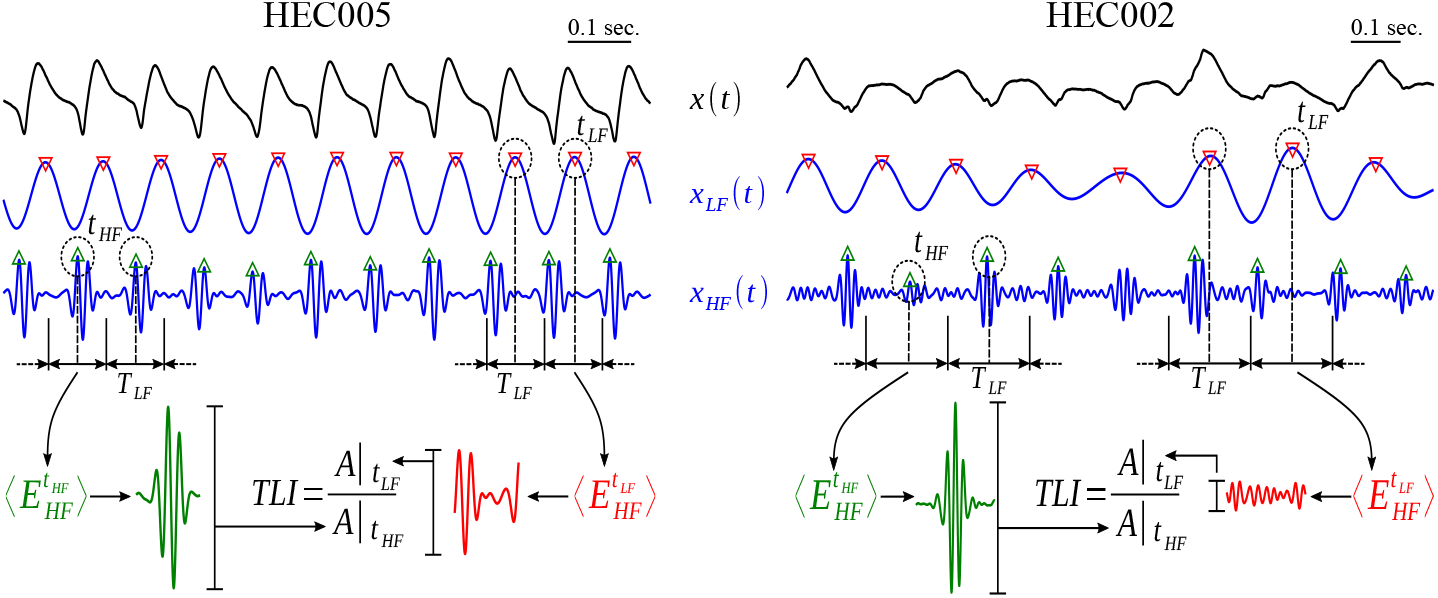
Ictal recordings (bipolar channels) obtained from the seizure onset zone (SOZ) and derived time series involved in the time locked index (TLI) algorithm for quantification of harmonicity. The low frequency (LF) band-pass filtered signal (*x_LF_* (*t*)) correspond to the Alpha and Theta bands for patients HEC005 and HEC002, respectively. The high frequency (HF) band-pass filtered signal (*x_HF_* (*t*)) correspond to the Gamma band for both patients. Using a time series of 7 sec. in length which included the epoch shown in these plots we obtained TLI = 0.57 and TLI = 0.15 for patients HEC005 and HEC002, respectively. Symbols and abbreviations: *x*(*t*), ictal LFP; *T_LF_*, period of the LF band; *t_LF_*, time instants corresponding to the maximum amplitude of the *x_LF_* time series; *t_HF_*, time instants corresponding to the maximum amplitude of the *x_HF_* time series; 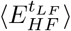, average of the epochs extracted from the *x_HF_* (*t*) time series with a length equal to *T_LF_* and centered at the *t_LF_* time instants; *A|t_LF_*, peak-to-peak amplitude of the mean epoch 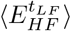; 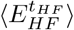, average of the epochs extracted from the *x_HF_* (*t*) time series with a length equal to *T_LF_* and centered at the *t_HF_* time instants; *A|t_HF_*, peak-to-peak amplitude of the mean epoch 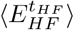.

In the case that the time series *x* was predominantly constituted by harmonic spectral components, the fast (*x_HF_*) and slow (*x_LF_*) oscillatory dynamics are characterized by a high degree of synchronization in time domain (i.e. phase-locking). As a consequence, the amplitude of 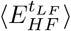 results comparable to that of the 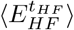 and so we obtain TLI *≈* 1 (see the case HEC005 in Figure 3). On the other hand, if the spectral energy of the time series *x* is not concentrated in narrow harmonically related frequency bands, the fast (*x_HF_*) and slow (*x_LF_*) rhythms will be not, in general, phase-locked. Therefore, the amplitude of 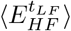 is averaged out to zero and TLI *≈* 0 is obtained for a sufficiently large number of samples *N_s_*. It is worth noting that the phase synchronization between the band-pass filtered time series (*x_LF_* and *x_HF_*) can be quantified using the PLV metric, however, the TLI algorithm has two significant advantages: 1) The TLI algorithm does not require to know the harmonic ratio between the two frequency bands of interest. In contrast, to compute the PLV you need to know this harmonic ratio, *a priori*, in order to be able to evaluate the phase-phase cross frequency coupling Tass et al. (1998). 2) The TLI metric can be effectively computed using slightly selective BPF, i.e. filters having wide bandwidths or low steepness of the transition bands. That is, by operating in the time-domain the TLI reliably assess the degree of time-locking, even in the case in which several (harmonic) spectral components are included within the bandwidth of the filters used to obtain the slow (*x_LF_*) and fast (*x_HF_*) rhythms. More details about the performance of TLI metric, its characterization using simulated data and links to the source code for TLI computation and test script examples can be found in Velarde et al. (2019).

### 2.7. Time series of PAC and harmonicity metrics

The TLI metric was tailored designed to be combined with the PC metric for improving the characterization and interpretation of the PAC patterns observed at the signal level. In line with this, time series were constructed for the PLV, KLMI, TLI and PC metrics to analyze their temporal evolution during the pre, post and ictal dynamics. Unless otherwise specified, the time series were constructed by computing all metrics in a sliding epoch of 5 sec. in length with 80 % overlap to include several periods for the slowest modulating rhythms explored (Delta band). This epoch length was an acceptable trade-off between statistical significance and temporal resolution capable to capture the PAC and harmonicity transients during the seizure dynamics. The ictal time series were computed spanning the entire duration of each seizure. Besides, the pre- and post-ictal time series were computed in time intervals of approximately 20 sec. in length located just before the seizure onset and just after the end of the seizure, respectively. In all cases, we use the onset and end of the seizure time marks determined by the epileptologists (SK and NC).

### 2.8. Distance matrix

To statistically quantify the extent of dependence between the PAC patterns and the harmonicity of the ictal activity, the distance matrix (DM) was computed for each patient as follows. The ictal time series (i.e. spanning the entire duration of each seizure) for each bipolar iEEG channel was constructed for the metrics of interest (KLMI, TLI) as described in Section 2.7. The resulting ictal time series corresponding to homologous iEEG channels were concatenated across all the seizures. Thus, we obtain a single time series per bipolar iEEG channel. Finally, the distance matrix was obtained by computing the Kolmogorov-Smirnov (K-S) statistic between all possible combinations of the time series taken in pairs, resulting in a matrix of size *N_c_ × N_c_* where *N_c_* is the number of bipolar iEEG channels.

### 2.9. Receiver operating characteristic analysis

The role of the harmonicity of the seizure dynamics in the performance of ictal PAC to identify the iEEG SOZ (seizure onset zone identified from the intracerebral electroencephalography recordings) compared to the classification given preoperatively by the epileptologists, was assessed by means of a receiver operating characteristic (ROC) analysis implemented as follows. Pre, post and ictal time series for each bipolar iEEG channel were constructed as described in Section 2.7 for each metric of interest (Power, PLV, KLMI, TLI) taking into account all the seizures associated to each patient (see Table 1). The Power time series were computed for all frequency bands listed in Table A.1 of Appendix A.1. Besides, the PAC (PLV, KLMI) and harmonicity (TLI) time series were computed for all suitable combinations between the modulating LF (High Delta to High Beta) and modulated HF (Alpha to High HFO, HFO: high frequency oscillations) frequency bands. The inclusion criteria consisted of: (I) the width of the modulated HF band must be at least two times the center frequency of the modulating LF band, which is about the minimum bandwidth required to detect PAC Berman et al. (2012), (II) the maximum frequency of the modulating LF band must be less than or equal to the minimum frequency of the modulated HF band, which prevents spurious PAC from occurring due to overlapping LF and HF frequency bands. We subsequently computed the mean values of the pre, post and ictal time series (Power, PLV, KLMI, TLI) for each bipolar iEEG channel, seizure and frequency band. These mean values assigned to the bipolar iEEG channels were used to compute the ROC curves. In the ROC plots, true positive rate (TPR) was plotted against false positive rate (FPR) for each suitable frequency band combination (modulating LF vs. modulated HF). TPR was defined as the proportion of bipolar iEEG channels pertaining to the SOZ meeting the moving threshold. FPR was defined as the proportion of bipolar iEEG channels not pertaining to the SOZ meeting the moving threshold. In defining the TPR and FPR, the classification of the iEEG channels involved in the SOZ given preoperatively by the epileptologists for each single seizure was taken as a ground truth. The overall power of iEEG SOZ classification for a given metric and frequency band was quantified as the area under the ROC curve (AUC), where AUC = 0 (TPR = FPR) implies random classification and AUC = 0.5 implies perfect classification.

### 2.10. Bivariate analysis

A bivariate analysis based on the covariance method was implemented to shed light on how the harmonicity of the seizure dynamics, as measured by the TLI metric, related the performance of ictal PAC to identify the iEEG SOZ compared to the classification given preoperatively by the epileptologists. Pre, post and ictal time series for each bipolar iEEG channel were constructed as described in Section 2.7 for harmonicity (TLI) and PAC metrics (PLV, KLMI) taking into account all the seizures associated to each patient (see Table 1). The PAC (PLV, KLMI) and harmonicity (TLI) time series were all computed for the same frequency band combination (modulating LF vs. modulated HF) that maximizes the PLV-based iEEG SOZ classification power rated by the AUC in each patient (see Table A.2). Next, the mean value of each time series was computed for subsequent analysis. As a result, we obtained one matrix of mean values *N_c_ ×N_sz_* per patient, time interval (Pre, post and ictal) and metric (PLV, KLMI, TLI), where *N_c_* is the number of bipolar iEEG channels and *N_sz_* is the number of seizures of the patient (see Table 1). Each matrix of mean values was Z-score normalized across iEEG channels. Then, the elements of the resulting matrix were segregated in two samples of Z-scored mean values (SOZ and non SOZ), following the iEEG SOZ classification given preoperatively by the neurologist. Each SOZ sample of Z-scored mean values for the PAC metrics (PLV, KLMI) was concatenated with the homologous SOZ sample of Z-scored mean values for the harmonicity metric (TLI) to obtain a SOZ sample matrix from which the SOZ sample covariance matrix was computed. The same procedure was followed to obtain the non SOZ sample covariance matrices. Then, The eigenvalue decomposition (EVD) of the SOZ and non SOZ sample covariance matrices was computed. As a result, this EVD allowed us to effectively quantify the distance between the SOZ and non SOZ samples as the Euclidean distance between the centroids of the covariance error ellipses in the PAC (PLV, KLMI) vs. harmonicity (TLI) subspace.

### 2.11. Statistical analysis

Statistical analyses where conducted in R R Core Team (2019). We found that most of the variables studied present a non-normal distribution (assessed via Kolmogorov-Smirnov and Anderson-Darling tests). Therefore, non-parametric statistical tests were used to evaluate significance. For the two groups analyses of non-paired samples, we use the permutation test based on random sampling without replacement (see Appendix A.2). In the case of paired samples we use the Wilcoxon signed rank test (see Appendix A.3). To assess the statistical significance of the PLV comodulograms, we compute a distribution of 1 × 10^3^ surrogate PLV values achieved by applying the PLV measure to sample shuffled *φ_aHF_* (*t*) time series. Assuming a normal distribution of the surrogate |PLV| values, a significance threshold (PLV_thresh_) is then calculated by using P *<* 0.001 after Bonferroni correction for multiple comparisons. A similar procedure was used to assess the statistical significance of the KLMI comodulograms (data not shown) and TLI harmonicity maps using sample shuffled *a_HF_* (*t*) and *x_HF_* (*t*) time series, respectively.

## 3. RESULTS

In this section we first describe a series of preliminary observations which motivated our working hypothesis about the relationship between spectral harmonicity and performance of ictal PAC patterns as predictors of the iEEG SOZ. Then, we present evidence supporting the hypothesis and, in the last part, we quantitatively analyze statistics derived from the recordings to validate it.

### 3.1. Harmonic and non harmonic PAC patterns observed during the seizure dynamics

Figure 1 shows ictal LFP recordings obtained from the iEEG SOZ of each patient (1 bipolar channel per patient). The insets show 2 sec. epochs to highlight the oscillatory waveform shapes and time scales during ictal events. In some patients, it is possible to identify a characteristic pulse shape or motif which is repeated in time giving rise to a (quasi)periodic non sinusoidal oscillatory pattern (e.g. HEC005 case in Figure 1). The Fourier decomposition of these non sinusoidal oscillatory rhythms is characterized by the presence of dependent, namely harmonically related, spectral components with their power prominently concentrated in narrow frequency bands (see Figures 4A, 4C and 4E). The presence of a (quasi)periodic monomorphic dynamics is less evident, or nearly absent, in the epochs corresponding to the patients HEC002 and HEC006 shown in Figure 1. In other cases, we observe a separation of time scales manifested as a slow non sinusoidal rhythm coupled to a fast oscillation far beyond spectral harmonics have decayed, in which the fundamental frequencies associated to each oscillatory dynamics seem to be independent (e.g. High Delta and HFO frequency bands in the case HRM011 shown in Figure 1). To examine the interplay between the different time scales observed in the LFP time series, a systematic exploration of the harmonicity and PAC patterns emerging during the seizure dynamics of the seven patients was implemented including all suitable combinations between the modulating LF (High Delta to High Beta) and modulated HF (Alpha to High HFO) frequency bands (see the inclusion criteria described in Section 2.9 and Table A.1). In Figure 2, the signals involved in the computation of the PLV and KLMI metrics for the quantification of PAC are shown, for ictal epochs corresponding to the patients HEC005 and HEC002. In both cases, it is possible to identify a degree of correlation between the phase time series of the LF component (*φ_LF_* (*t*)) and the amplitude envelope of the HF band (*a_HF_* (*t*)). These cross frequency couplings are evidenced by the phase-amplitude histograms shown in Figure 2, in which the mean amplitude of the fast oscillations depart from the uniform distribution revealing the existence of a preferred LF phase. The periodic monomorphic pattern constituting the ictal epoch of the patient HEC005 shown in Figures 1 and 2 strongly suggests that the observed PAC, as detected by the PLV and KLMI algorithms, is elicited by the presence of harmonic spectral components in the Fourier spectrum of the ictal epoch. In the spectrogram shown in Figure 4A (patient HEC005), it is possible to distinguish the fundamental component in, early in the seizure, the Alpha band and harmonic spectral components ranging from Beta to Low HFO band. This observation is supported by the spectrum shown in Figure 4C. On the other hand, this effect is less evident in the spectrum of the ictal activity for the HEC002 case, showing a fundamental component in the Theta band and less prominent spectral harmonics ranging from Alpha to possibly Low Gamma band (see Figure 4D). To differentiate these scenarios in a quantitative manner we introduce the TLI metric which exploit the fact that, for a fixed waveform, harmonically related frequency bands are intrinsically linked to phase locking oscillations in time domain. The computation of the TLI metric is illustrated in Figure 3 for the same ictal epochs shown in Figure 2. Figure 3 shows a more periodic pattern in time domain and a higher degree of synchronization between the fast and slow oscillations in the HEC005 case compared to the HEC002 case. These qualitative observations from the raw and band pass filtered signals was quantified by the TLI algorithm, which produced a higher TLI value for the HEC005 ictal epoch than the one obtained for the HEC002 case (see caption of Figure 3).

**Figure 4:**
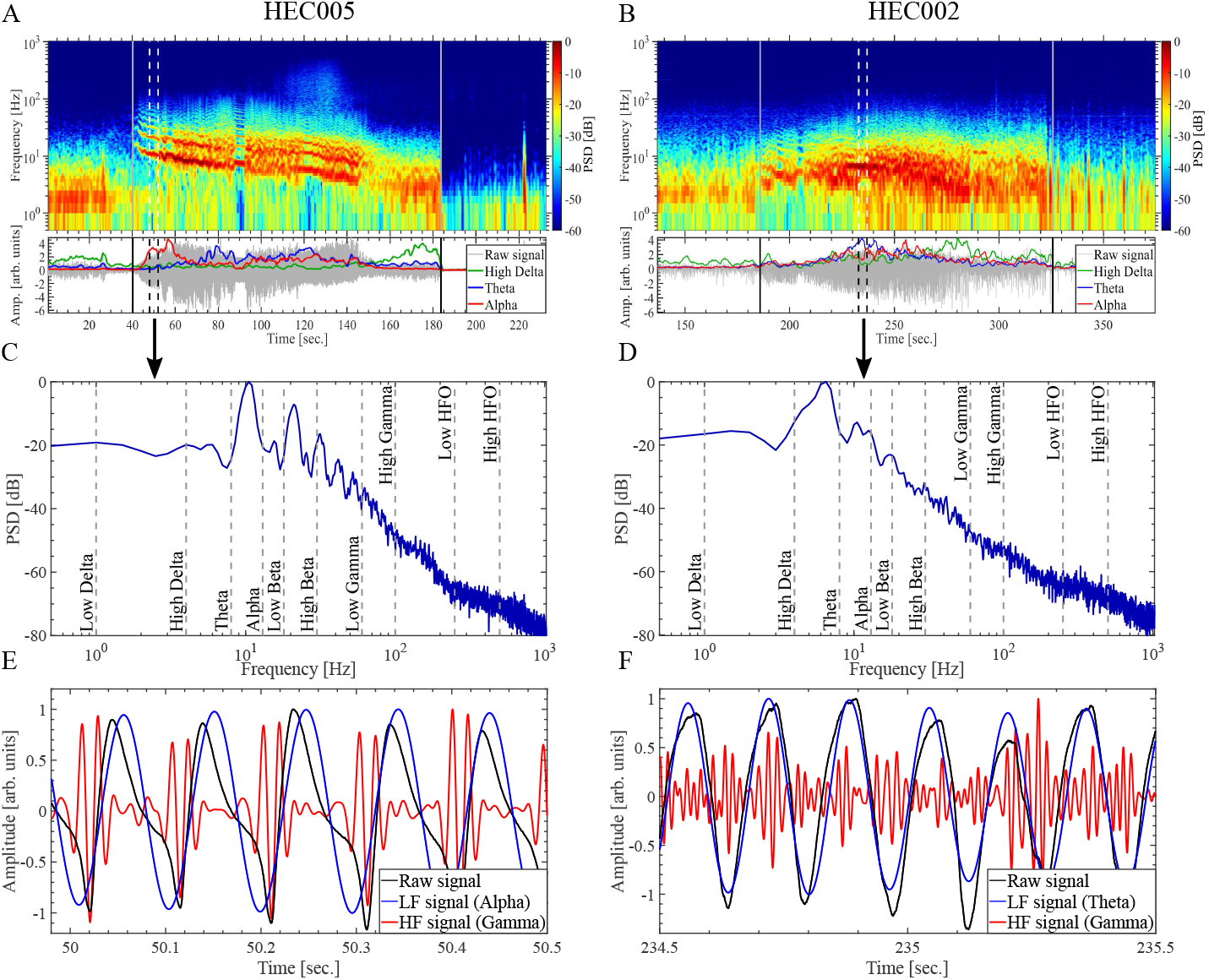
Spectrograms for the ictal activity of patients HEC005 and HEC002. (A, B) Fourier spectrograms computed using the Thomson multitaper method with a Hann window in the time domain. The time marks for the onset and termination of each seizure are indicated by the vertical solid gray lines. The lower panels show the ictal recordings (bipolar channels) obtained from the seizure onset zone (SOZ) of each patient (gray line), together with power time series for High Delta (green line), Theta (blue line) and Alpha (red line) frequency bands. Each power time series was Z-scored and smoothed by moving averaging technique to improve visualization (2 sec. sliding time window with 95 % overlap). The power time series were obtained by squaring the amplitude envelope of the signal band-pass filtered around each frequency band of interest. The amplitude envelopes were computed using the Filter-Hilbert method described in Section 2.5. (C, D) Power spectra of the bipolar recordings obtained from the time interval indicated by the vertical dashed lines in graphs at the top (panels A and B). (E, F) Bipolar recordings (black line) together with the low frequency (blue line) and high frequency (red line) band-pass filtered time series corresponding to the time interval indicated by the vertical dashed lines in graphs at the top (panels A and B). Symbols and abbreviations: HFO, high frequency oscillations; PSD, power spectral density; dB, decibel; HF, high frequency ; LF, low frequency.

### 3.2. Time locked plots

To examine the link between the LFP waveform shape and the types of PAC emerging during the seizure dynamics we can focus on the time locked plots shown in Figures 5, 6 and 7. The traces D were obtained by averaging approximately 20 epochs extracted from the raw signal A, of length equal to one period as defined by the LF band (signal B), centered at the time points corresponding to the peaks of the HF oscillation during each epoch (green up-pointing triangles in signal C). When this procedure does not average out then the LF and HF rhythms are coupled (see traces D in Figures 5, 6 and 7), that is, the maximum amplitude of the HF bursts (HF peaks) do not occur at random time instants but at a preferred phase of the LF rhythm. This PAC, was confirmed by computing the phase-amplitude histograms associated to the KLMI algorithm (data not shown). Importantly, the trace D in Figure 5 closely approximates the waveform observed as a repetitive pattern in the original raw signal (see Figure 5A) with no evident presence of a burst-like transient of Gamma oscillation. This observation suggests that the raw signal is constituted by other spectral components besides the Gamma oscillation, phase locked to the LF rhythm (i.e. harmonic spectral components) whose linear superposition delineates the non sinusoidal waveform shape of the periodic pattern. On the other hand, the trace D in Figure 7 shows the presence of a burst of Low HFO in a particular phase of the fundamental frequency defined by the non sinusoidal LF rhythm, suggesting that the slow and fast rhythms are likely independent. To analyze the phase locking between the oscillatory components constituting the raw signal we compute the TLP for the band-pass filtered signals. In Figures 5, 6 and 7, the traces E and F correspond to the time locked plots for the band-pass filtered signals shown in the graphs B and C, respectively. The trace E was obtained by averaging approximately 20 epochs extracted from the LF signal (graph B), of length equal to approx. 2 periods defined by the LF band, time locked to the LF peaks (red down-pointing triangles). The same procedure was used to obtain the signals shown in trace F, but in this case, the epochs time locked to the LF peaks were extracted from the HF signal (graph C). The low jitter level displayed by the time locked epochs shown in Figure 5E, highlights the periodicity of the fundamental rhythm constituting the ictal activity of patient HEC005. Moreover, the approximate coincidence of the average (black line) with any individual trace (gray line) in Figure 5F indicates that the Gamma oscillation is highly phase locked to the Alpha rhythm, that is, the fast and slow oscillations exhibit a harmonic frequency ratio. The Figure 7E shows a higher jitter level than that observed in Figure 5E indicating a relatively poorer periodicity for the High Delta rhythm corresponding to the HRM011 case. More important, the time locked epochs (gray line) displayed in Figure 7F show that the HF bursts occur around a particular phase of the LF rhythm indicating the existence of PAC, however, as the average (black line) of individual traces vanishes (see Figure 7F) the Low HFO oscillations are not phase locked to the High Delta rhythm. Taken together, these results show that in the HEC005 case the fast and slow oscillations are not independent but keep a harmonic frequency ratio with a fixed relative phase, which in turn is determined by the non sinusoidal waveform shape constituting the periodic ictal pattern. As a consequence, the coupling detected by the PLV and KLMI algorithm in this ictal scenario was termed as ‘harmonic PAC’. In contrast, the PAC observed in the HRM011 case results from independent, i.e. not time locked, slow and fast oscillations and it was termed ‘non harmonic PAC’. The traces E and F in Figure 6 corresponding to the patient HEC002 show an intermediate scenario with slightly lower periodicity of the LF rhythm and more attenuation of the time locked average obtained from the HF epochs (black line in Figure 6F) when compared to the HEC005 case (Figure 5F), suggesting less prominent harmonic components in the frequency domain for the HEC002 case (see also Figures 4B, 4D and 4F).

**Figure 5:**
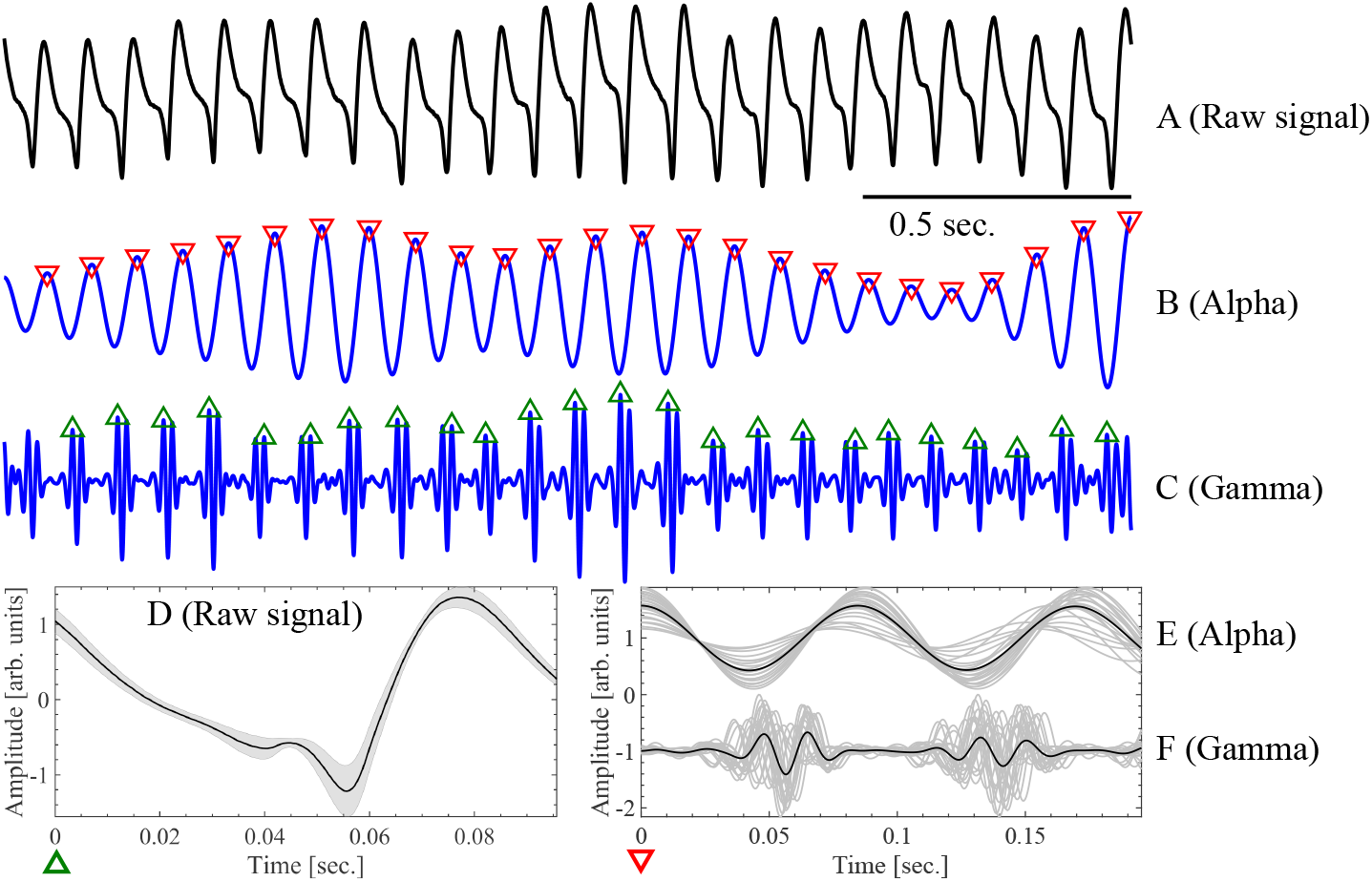
Time locked plots (TLP) computed from the ictal recordings (bipolar channels) obtained from the seizure onset zone (SOZ) of patient HEC005. (A) Unfiltered ictal recording (bipolar channel). (B) Ictal recording band-pass filtered around the Alpha frequency band. (C) Ictal recording band-pass filtered around the Gamma frequency band. (D), (E) and (F) are the TLP corresponding to the signals shown in graphs (A), (B) and (C), respectively.

**Figure 6:**
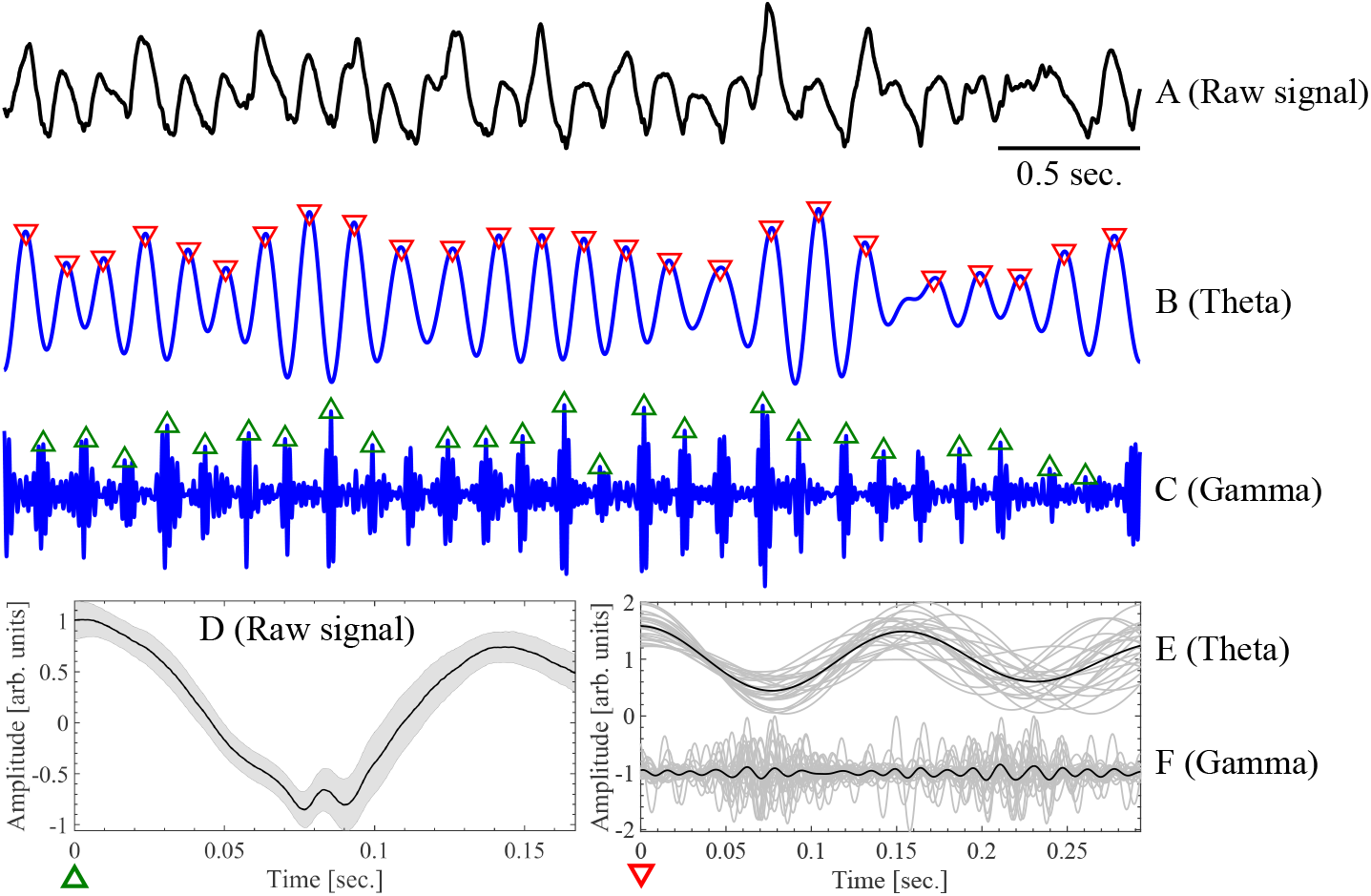
Time locked plots (TLP) computed from the ictal recordings (bipolar channels) obtained from the seizure onset zone (SOZ) of patient HEC002. (A) Unfiltered ictal recording (bipolar channel). (B) Ictal recording band-pass filtered around the Theta frequency band. (C) Ictal recording band-pass filtered around the Gamma frequency band. (D), (E) and (F) are the TLP corresponding to the signals shown in graphs (A), (B) and (C), respectively.

**Figure 7:**
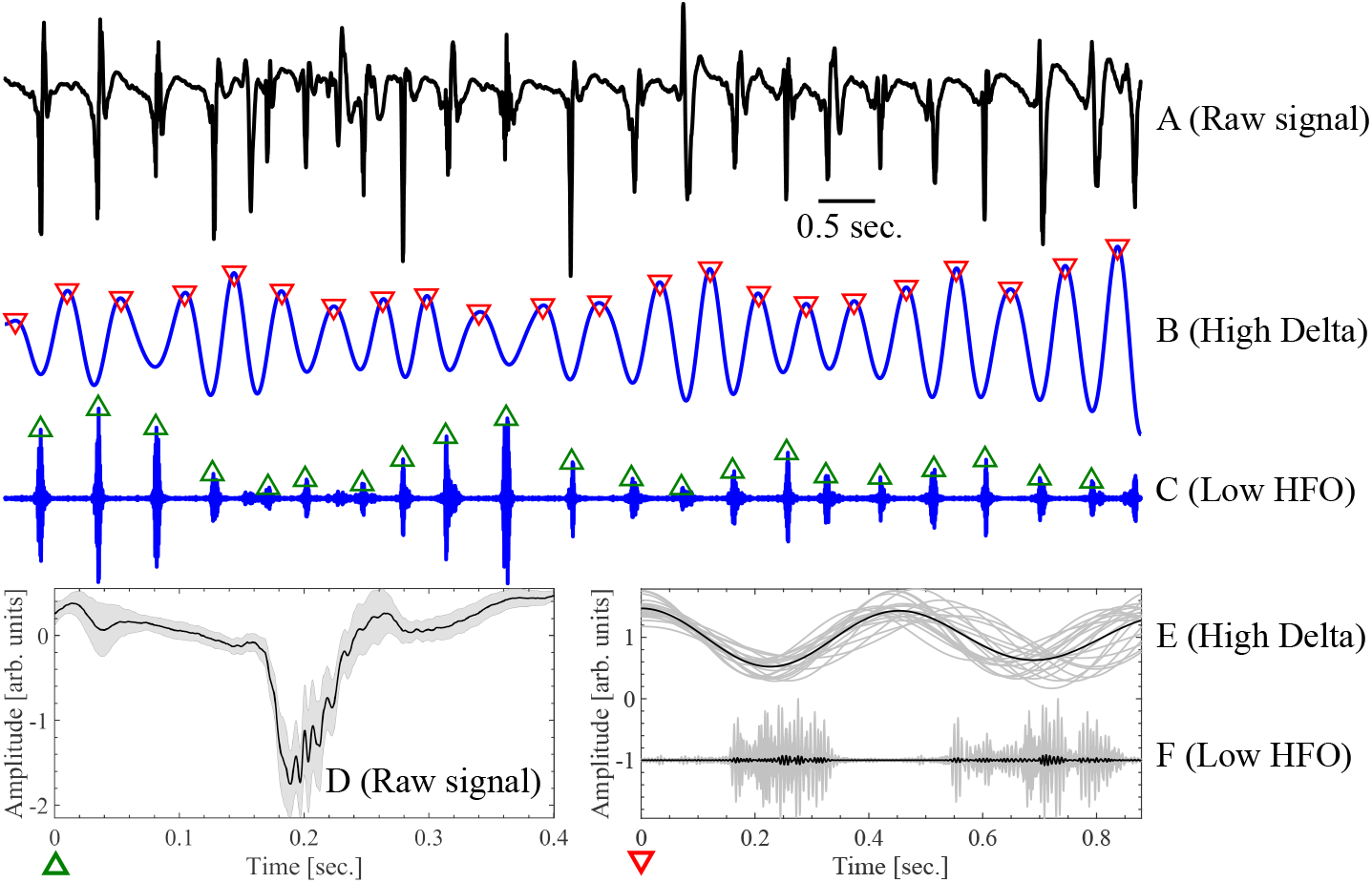
Time locked plots (TLP) computed from the ictal recordings (bipolar channels) obtained from the seizure onset zone (SOZ) of patient HRM011. (A) Unfiltered ictal recording (bipolar channel). (B) Ictal recording band-pass filtered around the High Delta frequency band. (C) Ictal recording band-pass filtered around the Low HFO frequency band. (D), (E) and (F) are the TLP corresponding to the signals shown in graphs (A), (B) and (C), respectively. Symbols and abbreviations: HFO, high frequency oscillations.

### 3.3. Comodulograms and harmonicity maps

To verify the consistency of the results obtained by examining the waveform shapes emerging during the ictal activity by means of the time locked plots (Figures 5, 6 and 7), we also analyzed the observed ictal scenarios in the frequency domain by computing the comodulograms and harmonicity maps shown in Figures 8 and 9. Figure 8 shows the PLV comodulograms and TLI harmonicity maps corresponding to ictal activity obtained from the iEEG SOZ for patients HEC005 and HEC002 (see Figures 1, 2, 3, 4, 5 and 6). Figures 8A and 8B show the PAC intensity as quantified by the PLV metric using a bandwidth of *Bw_HF_* = 50 Hz for the BPFs applied to extract the HF oscillations. Significant PAC was detected between Alpha vs. Gamma and Theta vs. Gamma frequency bands for the HEC005 and HEC002 cases, respectively. Importantly, the harmonicity map obtained for the HEC005 case (Figure 8C) shows significant TLI values co-occurrent in frequency with the PAC detected by the PLV metric (Figure 8A), further evidencing the presence of harmonic PAC between the Alpha vs. Gamma frequency bands during the seizure dynamics of patient HEC005. This conclusion is supported by the results shown in the panel E of Figure 8 corresponding to the PLV computed using narrow BPFs to extract the HF oscillations, i.e. setting *Bw_HF_* = 20 Hz which is close to the minimum bandwidth value required to detect PAC (two times the modulating frequency) Berman et al. (2012). This comodulogram highlights through the presence of multiple PAC intensity peaks related by rational factors, the fact that the PAC observed in the patient HEC005 is elicited by the presence of harmonic spectral components in the Fourier spectrum of the ictal activity. Again, this effect is less evident in the HEC002 case as it is shown in panels D and F of Figure 8. Figures 9C, 9D and 9E show the PLV comodulograms and TLI harmonicity maps corresponding to ictal activity obtained from the SOZ for the patient HRM011 (see Figures 1 and 7). The comodulogram shown in Figure 9C indicates the presence of significant PAC intensity between High Delta vs. High Gamma and HFO frequency bands. Moreover, the associated harmonicity map (Figure 9D) shows no significant TLI values associated to the observed PAC. The latter is consistent with the Fourier spectrogram and spectrum shown in Figures 9A and 9B, in which no evident harmonic spectral components pertaining to the HFO band are distinguishable. In fact, the spectrum in Figure 9A shows wide bumps in the High Gamma and HFO bands which have been associated to nested oscillations (i.e. non harmonic PAC) Lepage and Vijayan (2015). On the other hand, the comodulogram computed using narrow BPF to extract the HF oscillations (Figure 9E) highlights the presence of harmonic PAC between a modulating LF rhythm in the High Delta band (≈2 Hz) and the modulated rhythms in the Theta and Low Beta frequency bands (4 Hz - 20 Hz). This harmonic PAC is elicited by the harmonic spectral components, ranging from Theta to Beta bands, constituting the non sinusoidal ictal rhythm whose fundamental frequency lies in the High Delta band (see the spectrogram and the spectrum shown in Figures 9A and 9B and the waveform shapes shown in Figures 1, 7A and 7D).

**Figure 8:**
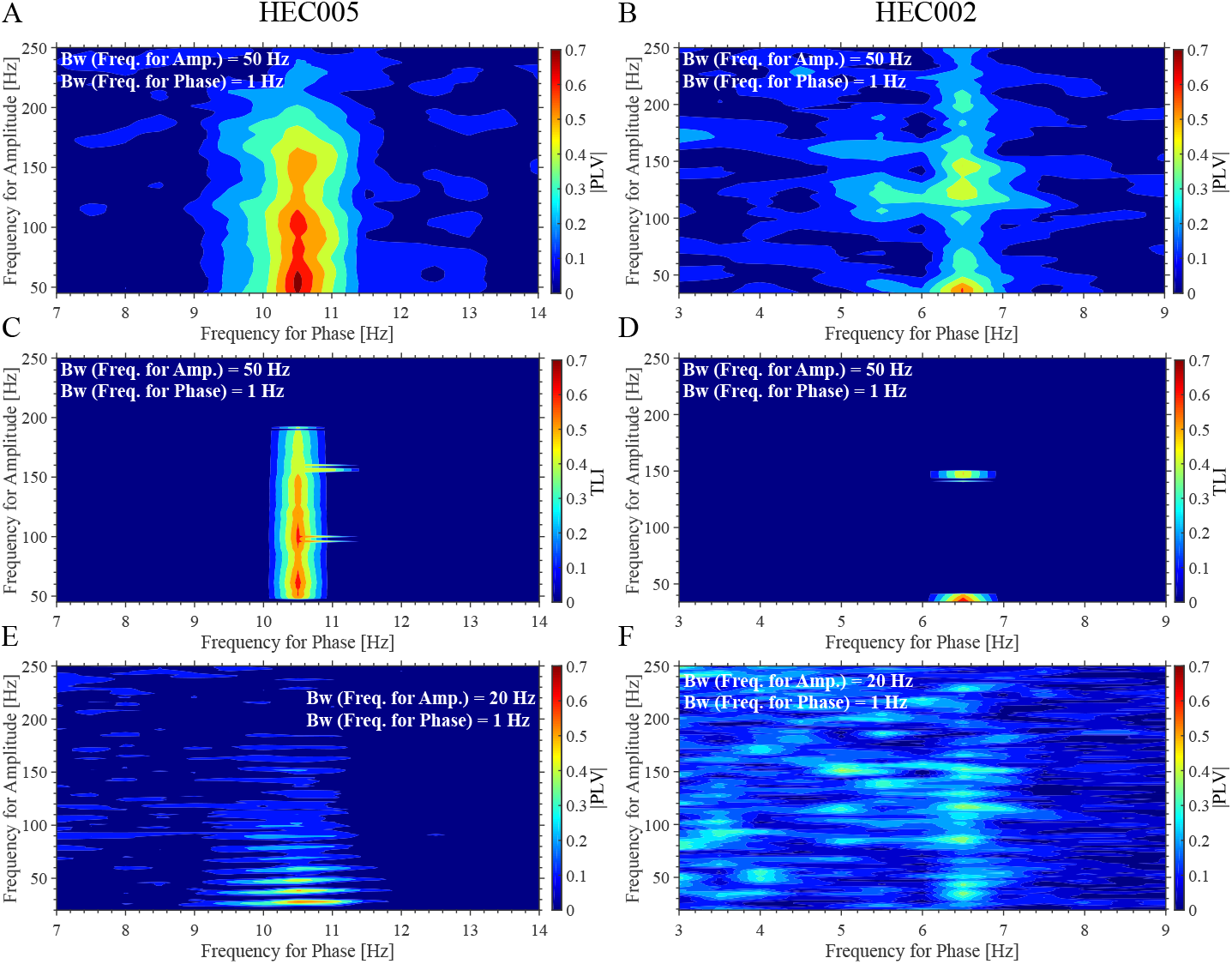
Phase-amplitude coupling (PAC) comodulograms and harmonicity maps for the ictal activity of patients HEC005 and HEC002. (A, B) PAC comodulograms computed using the phase locking value (PLV) metric on the ictal recordings (bipolar channels) obtained from the seizure onset zone (SOZ) of patients HEC005 and HEC002. Note that the bandwidth (*Bw_HF_*) of the band-pass filters (BPFs) used to obtain the high frequency (HF) oscillations (*Bw_HF_* = 50 Hz), is wider than two times the maximum modulating frequency (14 Hz in graph A and 9 Hz in graph B) Berman et al. (2012). (C, D) Time locked index (TLI) harmonicity maps computed using the same BPFs and from the same ictal activity corresponding to the comodulograms shown in graphs at the top (panels A and B). (E, F) PLV comodulograms computed from the same ictal activity corresponding to the comodulograms shown in graphs at the top (panels A and B). Note that in this case we use narrow BPFs to obtain the HF oscillations (*Bw_HF_* = 20 Hz). The statistical significance of PLV comodulograms and time TLI maps was assessed as described in Section 2.11. Symbols and abbreviations: Bw (Freq. for Amp.) = *Bw_HF_*, bandwidth of the band-pass filters used to obtain the frequency for amplitude (HF oscillations); Bw (Freq. for Phase) = *Bw_LF_*, bandwidth of the band-pass filters used to obtain the frequency for phase (low frequency (LF) oscillations).

**Figure 9:**
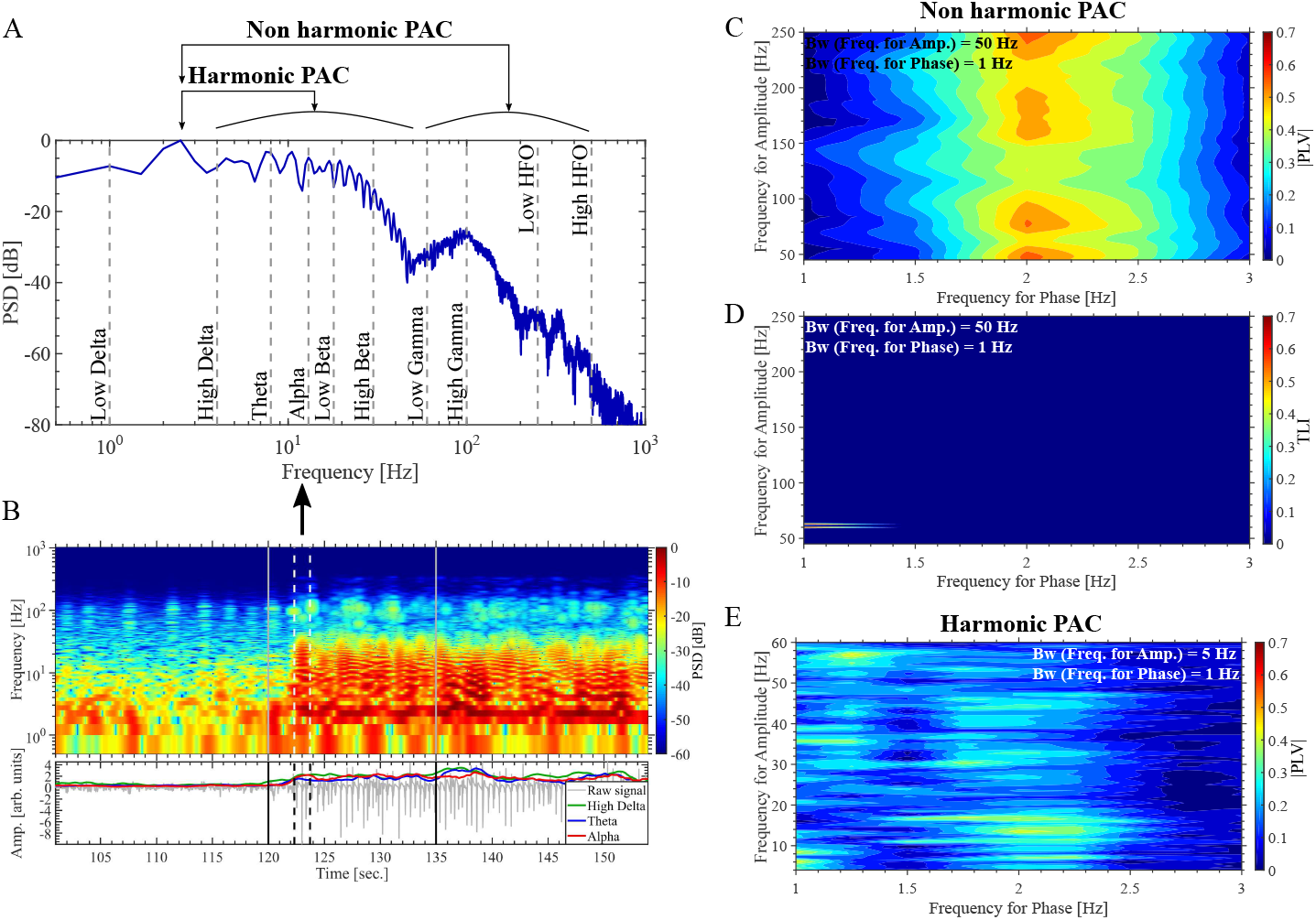
Spectrograms, phase-amplitude coupling (PAC) comodulograms and harmonicity maps for the ictal recordings (bipolar channels) obtained from the seizure onset zone (SOZ) of patient HRM011. (A) Power spectra of the ictal recording obtained from the time interval indicated by the vertical dashed lines in the Fourier spectrogram shown in panel B. (B) Fourier spectrogram computed using the modified periodogram method with a Hann window in the time domain. The time marks for the onset and termination of each seizure are indicated by the vertical solid gray lines. The lower panel show the ictal recording (bipolar channel) obtained from the SOZ of patient HRM011, together with power time series for High Delta, Alpha and Theta frequency bands. Each power time series was Z-scored and smoothed by moving averaging technique to improve visualization (2 sec. sliding time window with 95 % overlap). The power time series were obtained by squaring the amplitude envelope of the signal band-pass filtered around each frequency band of interest. The amplitude envelopes were computed using the Filter-Hilbert method described in Section 2.5. (C) PAC comodulograms computed using the phase locking value (PLV) metric on the ictal recording obtained from the SOZ of patient HRM011. This comodulogram was computed using wide band-pass filters (BPFs) to extract the high frequency (HF) oscillations (*Bw_HF_* = 50 Hz). (D) Time locked index (TLI) harmonicity map computed using the same BPFs and from the same ictal activity corresponding to the comodulogram shown in panel C. (E) PLV comodulogram computed from the same ictal activity corresponding to the comodulogram shown in panel C. Note that in this case we use narrow BPFs to extract the HF oscillations (*Bw_HF_* = 5 Hz). The statistical significance of PLV comodulograms and TLI maps was assessed as described in Section 2.11. Symbols and abbreviations: PSD, power spectral density; dB, decibel; Bw (Freq. for Amp.) = *Bw_HF_*, bandwidth of the band-pass filters used to obtain the frequency for amplitude (HF oscillations); Bw (Freq. for Phase) = *Bw_LF_*, bandwidth of the band-pass filters used to obtain the frequency for phase (low frequency (LF) oscillations).

### 3.4. Temporal evolution of PAC and harmonicity metrics during the seizure dynamics

The connection between PLV and TLI metrics associated to the emergence of harmonic and non harmonic PAC patterns was assessed by analyzing their temporal evolution across bipolar iEEG channels. Figures 10 and 11 show the temporal evolution of the PLV, TLI and PC metrics during a single seizure for patients HEC005 and HEC002, respectively. In Figures 10 and 11 the PLV, TLI and PC time series were constructed as described in Section 2.7. In both patients, it is possible to distinguish that the PLV intensity (solid black line) increases close to the seizure onset in some channels pertaining to the iEEG SOZ. The propagation profile of the PAC pattern (dotted green line) was determined from the time instant in each channel (green filled circles) at which the PLV intensity exceeds the threshold given by 3 times its median value computed from the pre-ictal time interval. Due to the fact that we used sufficiently narrow BPF to obtain an almost sinusoidal low frequency component (i.e. uniform distribution of the phase values in *φ_LF_* (*t*), see Section 2.5), the PC time series (blue solid line) is close to PC ≈ 0 along the seizure dynamics in all the bipolar iEEG channels shown in Figures 10 and 11. This indicates that the observed PAC patterns are not spurious artifacts related to the presence of phase clustering in the modulating LF component Cohen (2008, 2014). In connection with Figures 1, 10, 11, 12 and 13A, it is important to note a key point regarding the interpretation of TLI as a measure complementary to the estimators aimed to quantify PAC (e.g. PLV and KLMI). Even though the TLI metric is bounded in the range [0, 1] (see Section 2.6), the mean value of the TLI time series does depend on the noise level present in the processed LFP and on the epoch length, i.e. the number of periods of the low frequency oscillation taken to implement the time-locked average involved in the TLI computation (see Figure 13A). As a consequence, a robust detection of harmonic PAC should be based on observing the evolution of PAC and TLI metrics as functions of time (or frequency as shown in Figures 8 and 9, iEEG channels or any other parameter of interest) and not on a direct comparison against an absolute magnitude threshold. Therefore, the key point indicating the presence of harmonic PAC is given by the fact that the TLI increases concurrently with the PAC metrics (PLV and KLMI), rather than by the absolute TLI value at a particular time instant. In this regard, Figure 10 shows the PAC and harmonicity patterns for the patient HEC005 in which the intensity of the PLV and TLI time series increase simultaneously being this behavior a robust indicator about the presence of harmonic PAC. For the patient HEC002, Figure 11 shows that the PLV intensity also increases close to the seizure onset for the channels pertaining to the iEEG SOZ, although in this case the simultaneous increase of the TLI intensity is highly attenuated. The potential of the TLI and PC metrics as tools complementary to the PAC metrics (PLV, KLMI) for the interpretation of PAC patterns is illustrated in Figure 1 where the temporal evolution of these metrics can be contrasted to the waveform shapes emerging in the ictal scenarios, for the seven patients included in this study. In particular, the non harmonic PAC pattern between High Delta vs. HFO bands for the patient HRM011 is evidenced by the increase of the PLV intensity at the seizure onset without a concurrent increase of the TLI intensity.

**Figure 10:**
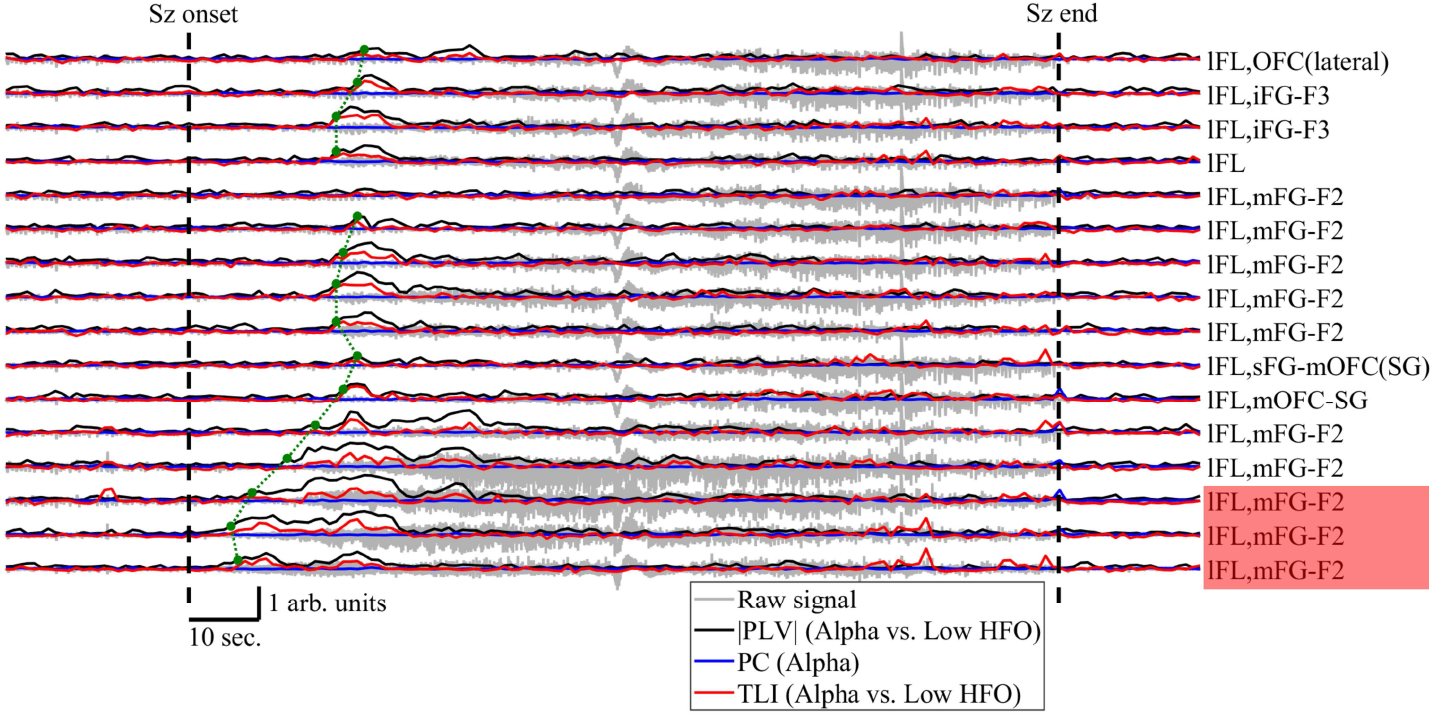
Temporal evolution of the recordings (bipolar channels, gray line) together with phase-amplitude coupling (PAC) and harmonicity metrics for the patient HEC005. The time marks for the onset and termination of each seizure (Sz) are indicated by the vertical dashed black lines. The Phase locking value (PLV, black line) quantifying PAC, time locked index (TLI, red line) quantifying harmonicity and Phase clustering (PC, blue line) time series were constructed using a sliding epoch of 2 sec. in length with 50 % overlap (see Section 2.7). In all the cases, the offset of the TLI time series was removed by substracting the mean value of the pre-ictal TLI time series computed from a time interval of approximately 20 sec. in length located just before the seizure onset. The propagation profile of PAC (dashed green line) was determined from the time instant in each bipolar channel (green dots) at which the PLV time series exceeds the threshold given by 3 times its median value computed from the pre-ictal time interval. The channels highlighted in red correspond to the seizure onset zone (SOZ). The labels of bipolar channels of patient HEC005 are described in Table A.3. Symbols and abbreviations: HFO, high frequency oscillations.

**Figure 11:**
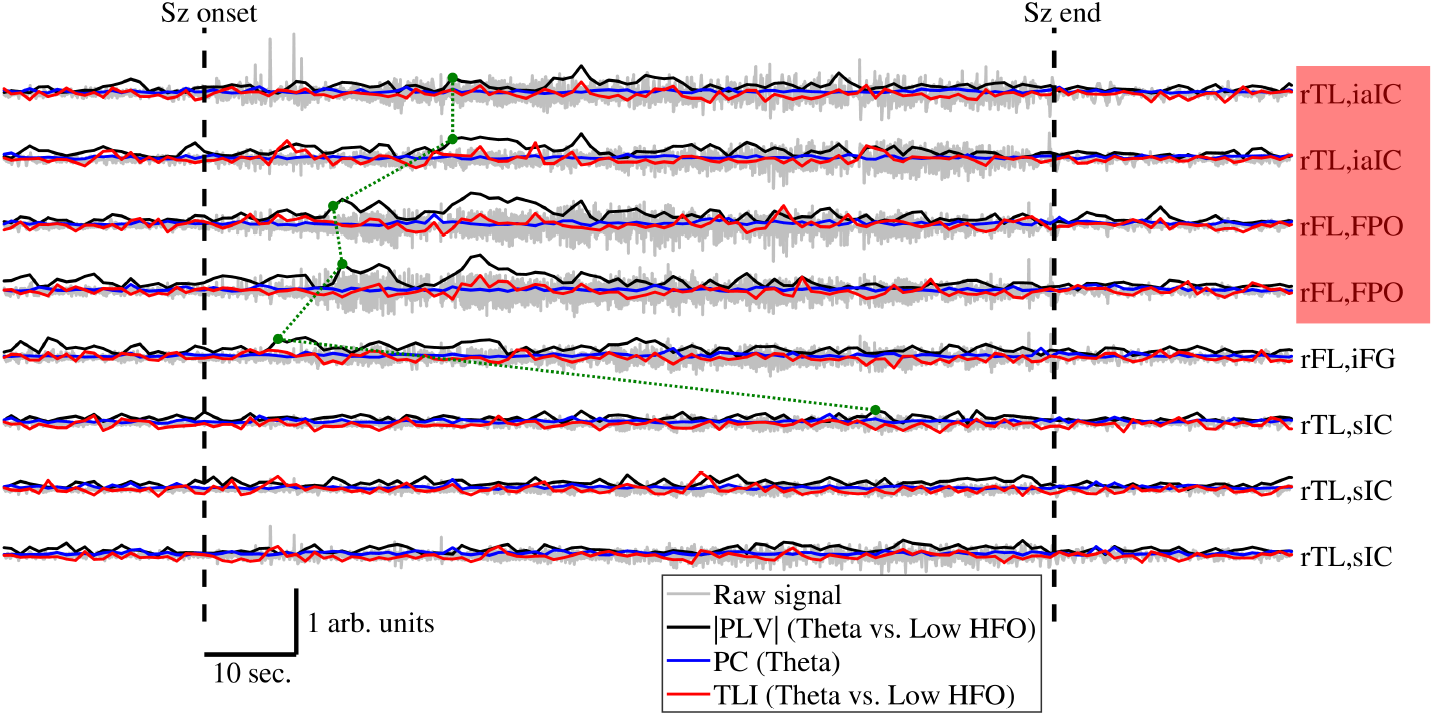
Temporal evolution of the recordings (bipolar channels, gray line) together with phase-amplitude coupling (PAC) and harmonicity metrics for the patient HEC002. The time marks for the onset and termination of each seizure (Sz) are indicated by the vertical dashed black lines. The Phase locking value (PLV, black line) quantifying PAC, time locked index (TLI, red line) quantifying harmonicity and Phase clustering (PC, blue line) time series were constructed using a sliding epoch of 2 sec. in length with 50 % overlap (see Section 2.7). In all the cases, the offset of the TLI time series was removed by substracting the mean value of the pre-ictal TLI time series computed from a time interval of approximately 20 sec. in length located just before the seizure onset. The propagation profile of PAC (dashed green line) was determined from the time instant in each channel (green dots) at which the PLV time series exceeds the threshold given by 3 times its median value computed from the pre-ictal time interval. The channels highlighted in red correspond to the seizure onset zone (SOZ). The labels of bipolar channels of patient HEC002 are described in Table A.4. Symbols and abbreviations: HFO, high frequency oscillations.

**Figure 12:**
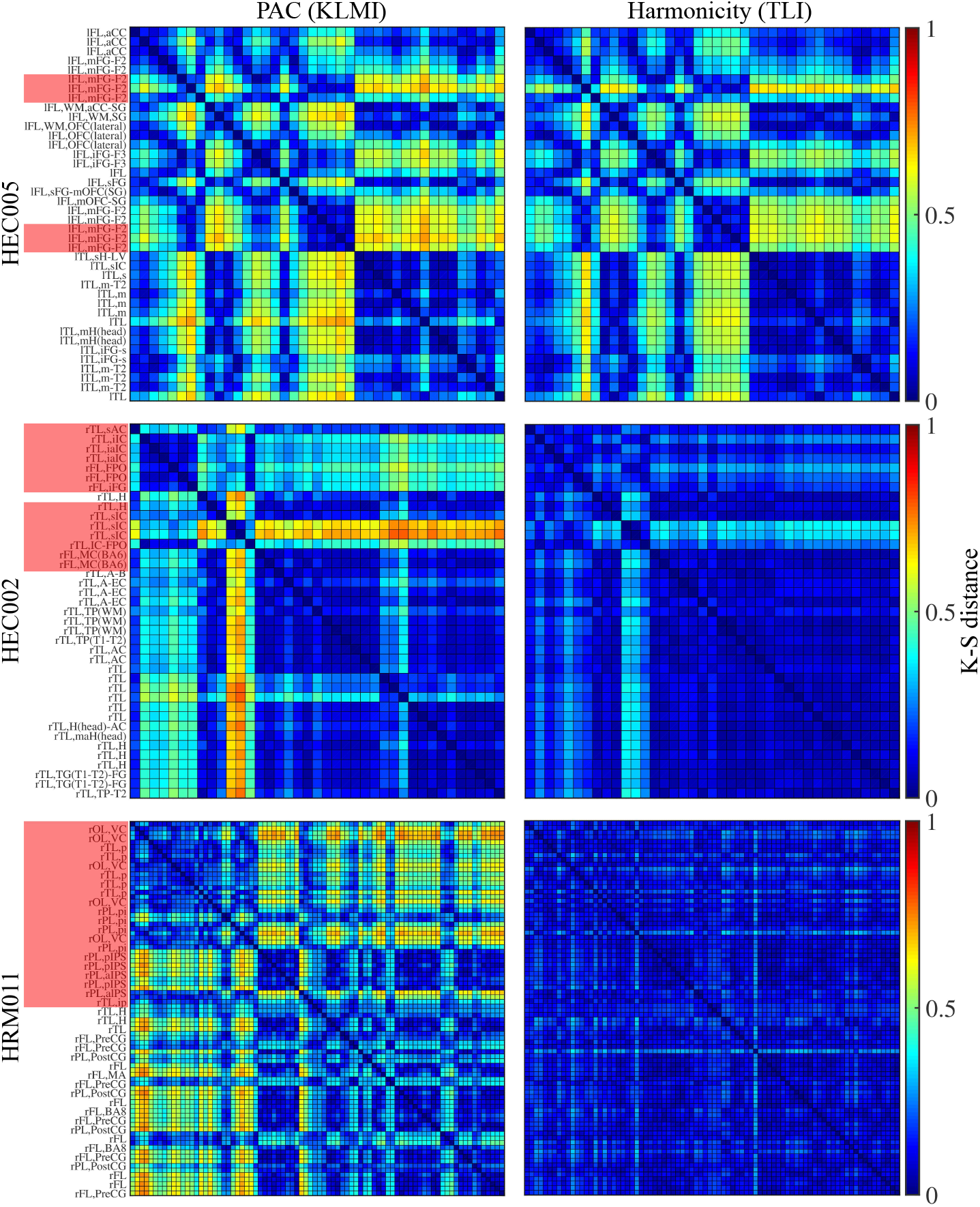
Kolmogorov-Smirnov (K-S) distance matrices for phase-amplitude coupling (PAC) and harmonicity (see Section 2.8). The distance matrices corresponding to the PAC (KLMI, modulation index based on the Kullback-Leibler distance) and harmonicity (TLI, time locked index) metrics were computed during the ictal time interval (i.e. spanning the entire duration of each seizure) and using the following frequency bands in each patient. HEC005: Beta vs. Gamma (40 bipolar channels); HEC002: Theta vs. Gamma (39 bipolar channels); HRM011: High Delta vs. HFO (82 bipolar channels. For clarity, labels are shown for every other bipolar channel). The channels highlighted in red correspond to the seizure onset zone (SOZ). The labels of bipolar channels of patients HEC005, HEC002 and HRM011 are described in Tables A.3, A.4 and A.5, respectively.

**Figure 13:**
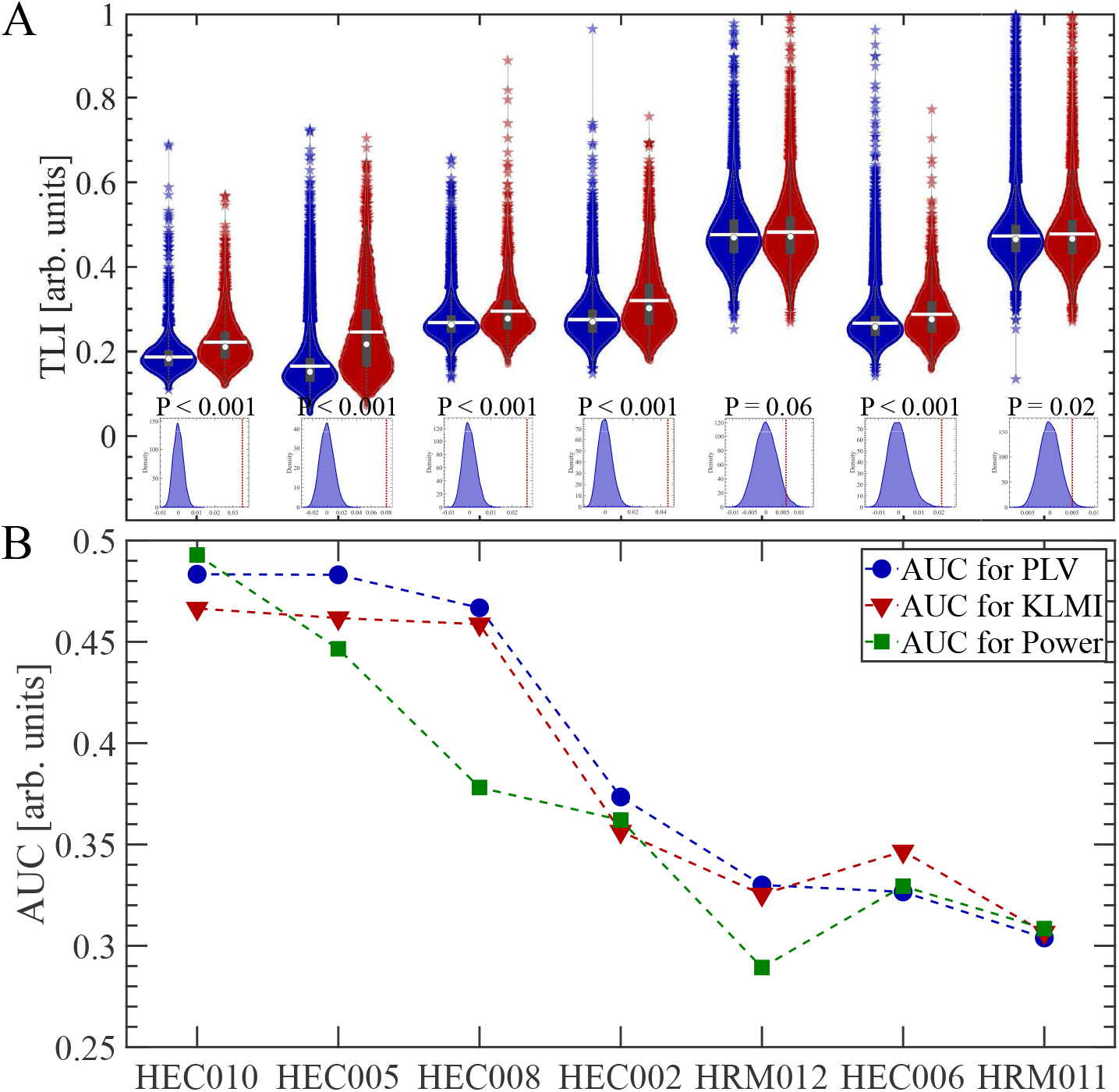
Receiver operating characteristic (ROC) analysis implemented for the ictal time interval. (A) Red and blue violin plots correspond to the seizure onset zone (SOZ) and non SOZ sample distributions of time locked index (TLI) values, respectively (see description of the two group analysis in Appendix A.2). Center gray boxes represent the 25th and 75th percentiles, whiskers (doted gray lines) extend to the most extreme data points not considered outliers (1.5 times the interquartile range), star markers represent outliers. The center white circle and white line indicate the median and mean, respectively. The histograms below the violin plots, correspond to the sampling distribution of the difference between the means of the SOZ and non SOZ distributions of TLI values. The histograms were computed via random sampling without replacement (1 10^4^ permutations). The vertical doted line (red) in the histograms indicates the actual mean difference between the SOZ and non SOZ distributions of TLI values shown in the corresponding violin plots. (B) Maximum AUC values corresponding to the ROC analysis based on the spectral power (Power) and PAC (PLV, KLMI) metrics (see Table A.2). The AUC values corresponding to the PAC (PLV, KLMI) metrics were computed for the frequency band combination (modulating LF vs. modulated HF) that maximizes the PLV-based SOZ classification power rated by the AUC in each patient (see Table A.2). Symbols and abbreviations: PAC, phase-amplitude coupling; PLV, phase locking value; KLMI, modulation index based on the Kullback-Leibler distance; AUC, area under the ROC curve.

### 3.5. Distance matrices

In this section, we started by analyzing the waveform shapes associated to epochs localized in brief time intervals during the ictal activity. Then, we discussed the dynamics of PAC and its complementary metrics across some iEEG channels during a single seizure. Following this bottom-up approach, Figure 12 shows distance matrices (DMs) for the KLMI and TLI metrics to illustrates the extent of dependence between PAC and harmonicity tacking into account all the iEEG channels and seizures of a given patient (see Section 2.8). It was found that the PAC and harmonicity patterns emerging across bipolar iEEG channels in the DMs, reproduce the results discussed above in terms of ictal waveform shapes and PAC temporal patterns. Specifically, the DMs for the patient HEC005 show that the bipolar iEEG channels presenting a high K-S distance for the KLMI values (Figure 12 upper-left), show a similar behavior for the TLI metric (Figure 12 upper-right), and thus, suggesting a high level of dependence between PAC and harmonicity during the ictal activity in this patient. This behavior is also observed, although in a less extent, in the patient HEC002 (Figure 12 middle-left and middle-right). In contrast, the KLMI pattern observed across the bipolar iEEG channels of subject HRM011 is not reproduced by the DM corresponding to the TLI metric (Figure 12 bottom-left and bottom-right), supporting the results discussed above showing the presence of non harmonic ictal PAC in this patient.

### 3.6. Two group and paired analyses

The results corresponding to the two group and paired analyses for the PAC and harmonicity metrics are described and discussed in Appendix A.2 and Appendix A.3 in connection with Figures A.1, A.2, A.3 and A.4.

### 3.7. Harmonicity of the seizure activity determines the performance of the ictal PAC as a biomarker to identify the iEEG SOZ

Following the results presented in the previous section, we next asked how the performance of ictal PAC to identify iEEG SOZ compared to the classification given preoperatively by the epileptologists was related to the harmonicity observed in the seizure dynamics itself. To this purpose, we implemented a receiver operating characteristic (ROC) analysis (see Section 2.9). A detailed discussion on the ROC curves obtained for the patients HEC005, HEC002 and HRM011 is given in Appendix A.4 in connection with Figures A.5, A.6 and A.7). Figure 13B shows the maximum AUC values corresponding to the ROC analysis based on the spectral power (Power) and PAC (PLV, KLMI) metrics, obtained for the seven patients. The frequency bands corresponding to the maximum AUC for each patient are listed in Table A.2. It was found that the PAC metrics exhibit a performance comparable or higher than that of the Power metric to reproduce the iEEG SOZ classification determined preoperative by the epileptologist (see Figure 13B and Table A.2). Figure 13A shows the sample distributions of TLI values computed for the frequency band combination (modulating LF vs. modulated HF) that maximizes the PLV-based iEEG SOZ classification power rated by the AUC in each patient (see Table A.2). The SOZ and non SOZ sample distributions of TLI values include all bipolar iEEG channels and seizures in each patient and are represented as red and blue violin plots, respectively (see description of the two group analysis in Appendix A.2). Importantly, this analysis suggests a degree of correlation between the AUC values for PAC metrics (PLV, KLMI) and the harmonicity of ictal activity in terms of both I) the difference between the mean values of the TLI distributions corresponding to the SOZ and non SOZ groups (see Figures 13) and II) the AUC values for TLI (see Figure A.8).

### 3.8. Inter-patient bivariate analysis

The findings discussed above motivated us to implement a bivariate analysis (see Section 2.10) designed for an in-depth examination on how the harmonicity of the seizure dynamics, as measured by the TLI metric, match the performance of ictal PAC to identify the iEEG SOZ compared to the classification given preoperatively by the epileptologists. Figures 14B and 14C show the SOZ (red filled circles) and non SOZ (blue filled circles) samples of Z-scored mean values in the PAC vs. harmonicity plane (PLV, TLI) for the patients HEC005 and HRM011, which were computed for the ictal time interval and the frequency band combination (modulating LF vs. modulated HF) that maximizes the PLV-based iEEG SOZ classification power rated by the AUC in each patient (see Table A.2). Figure 14B shows that the separation between the SOZ and non SOZ samples (solid black line connecting the centroids of the ellipses) occurs along the diagonal, supporting the evidence discussed above with regard to the presence of harmonic PAC in the ictal activity of patient HEC005. Besides, Figure 14C shows that the separation between the SOZ and non SOZ samples computed from the ictal time interval is much more higher along the PLV-axis than along the TLI-axis in the PAC vs. harmonicity plane. The latter, is consistent with the presence of High Delta vs. HFO non harmonic PAC in the ictal activity of patient HRM011 as it was discussed above (see also the results for the patient HRM011 in Appendix A.4). Figure 14A shows the Euclidean distance between the centroids of the covariance error ellipses in the PAC vs. harmonicity plane (PLV, TLI) computed from the ictal time interval for the seven patients. In Figure 14A, the black cross in the origin and the colored markers represents the centroids of the non SOZ and SOZ covariance error ellipses, respectively. It is worth noting that Figures 14B and 14C and the filled markers in Figure 14A were computed including all the seizures of each patient at once, as described in Section 2.10. On the other hand, the empty markers in Figure 14 were obtained by applying the procedure detailed in Section 2.10 over each seizure individually. That is, we obtained an Euclidean distance between the SOZ and non SOZ samples in the PAC vs. harmonicity plane (PLV, TLI) for each single seizure of a given patient. Importantly, the markers shown in Figure 14A are organized around a pattern which extents along the diagonal of the PAC vs. harmonicity plane (PLV, TLI). The latter indicates that the increase of Euclidean distance between the centroids of the covariance error ellipses, indicative of a better putative classification, is accompanied by an increase in the harmonicity of the PAC pattern as measured by the TLI metric. Thus, in considering the seven patients analyzed in this study the harmonic ictal PAC performs better than the non harmonic ictal PAC to identify the iEEG SOZ, compared to the classification given preoperatively by the epileptologists (see the AUC scale in the colorbar of Figure 14A). In addition, it was found that the inter-patient behavior observed in Figure 14A is independent of the metric used to quantify PAC (see the discussion about Figure A.9 in Appendix A.5). Moreover, Figure 15 shows the results of the bivariate analysis for pre, post and ictal time intervals indicating that the relation between the performance of PAC to identify the iEEG SOZ and the harmonicity is specific of the ictal time interval.

**Figure 14:**
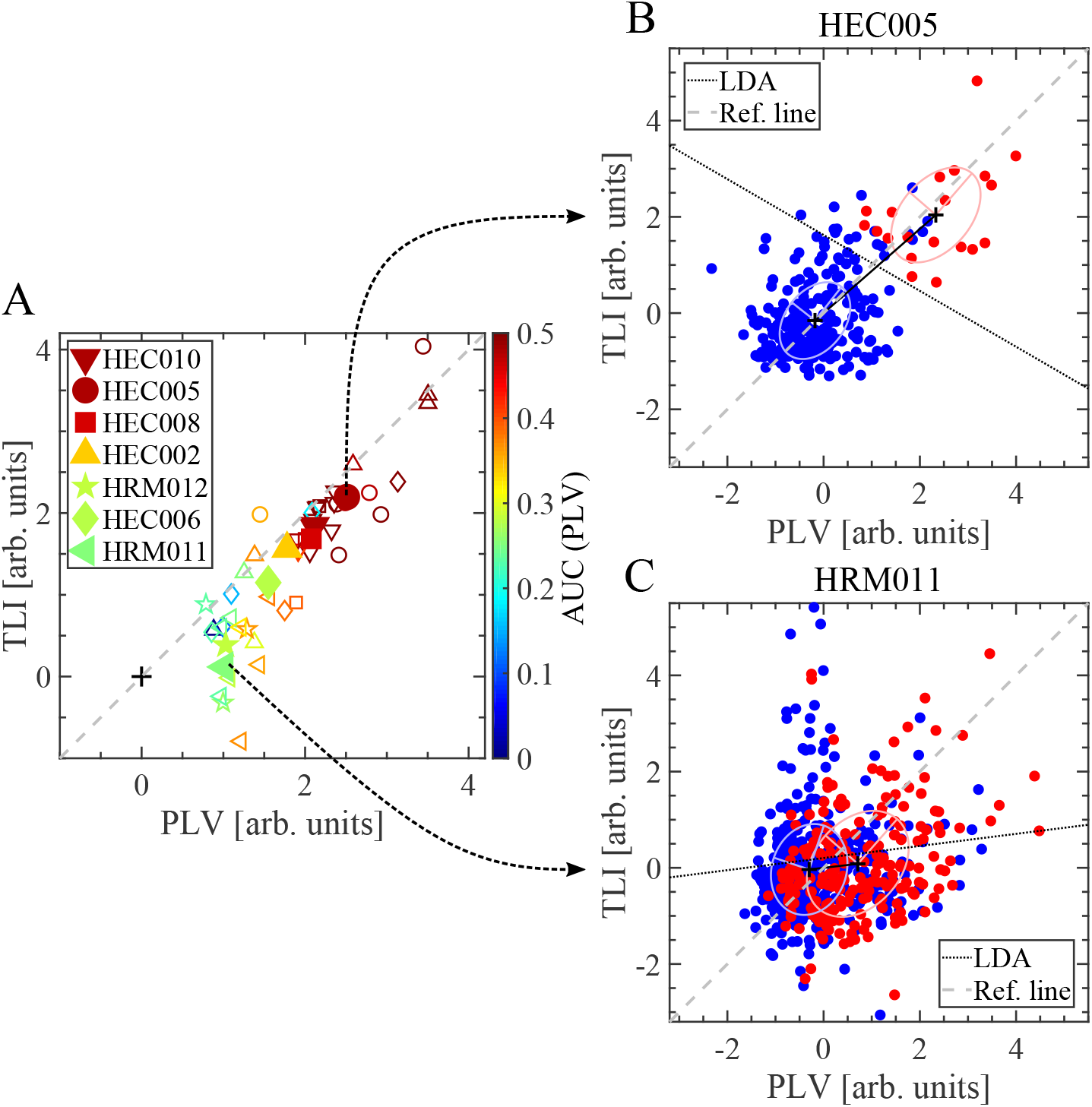
Bivariate analysis implemented for the ictal time interval (i.e. spanning the entire duration of each seizure). (A) Euclidean distance between the centroids of the covariance error ellipses. The black cross in the origin and the colored markers represents the centroids of the non SOZ and SOZ covariance error ellipses, respectively. Filled markers were computed including all the seizures of each patient at once. On the other hand, the empty markers were computed for a given patient taking each seizure individually. (B, C) Seizure onset zone (SOZ, red filled circles) and non SOZ (blue filled circles) samples of Z-scored mean values in the PAC vs. harmonicity plane (PLV, TLI) including all the seizures of each patient. Eigenvectors are shown together with the covariance error ellipses. The doted black line correspond to the Fisher’s discriminant resulting from the linear discriminant analysis (LDA) and the dashed gray line is the reference line. These graphs were computed for the frequency band combination (modulating LF vs. modulated HF) that maximizes the PLV-based SOZ classification power rated by the AUC in each patient (see Table A.2). Symbols and abbreviations: LF, low frequency; HF, high frequency; PAC, phase-amplitude coupling; PLV, phase locking value; TLI, time locked index; ROC, receiver operating characteristic; AUC, area under the ROC curve.

**Figure 15:**
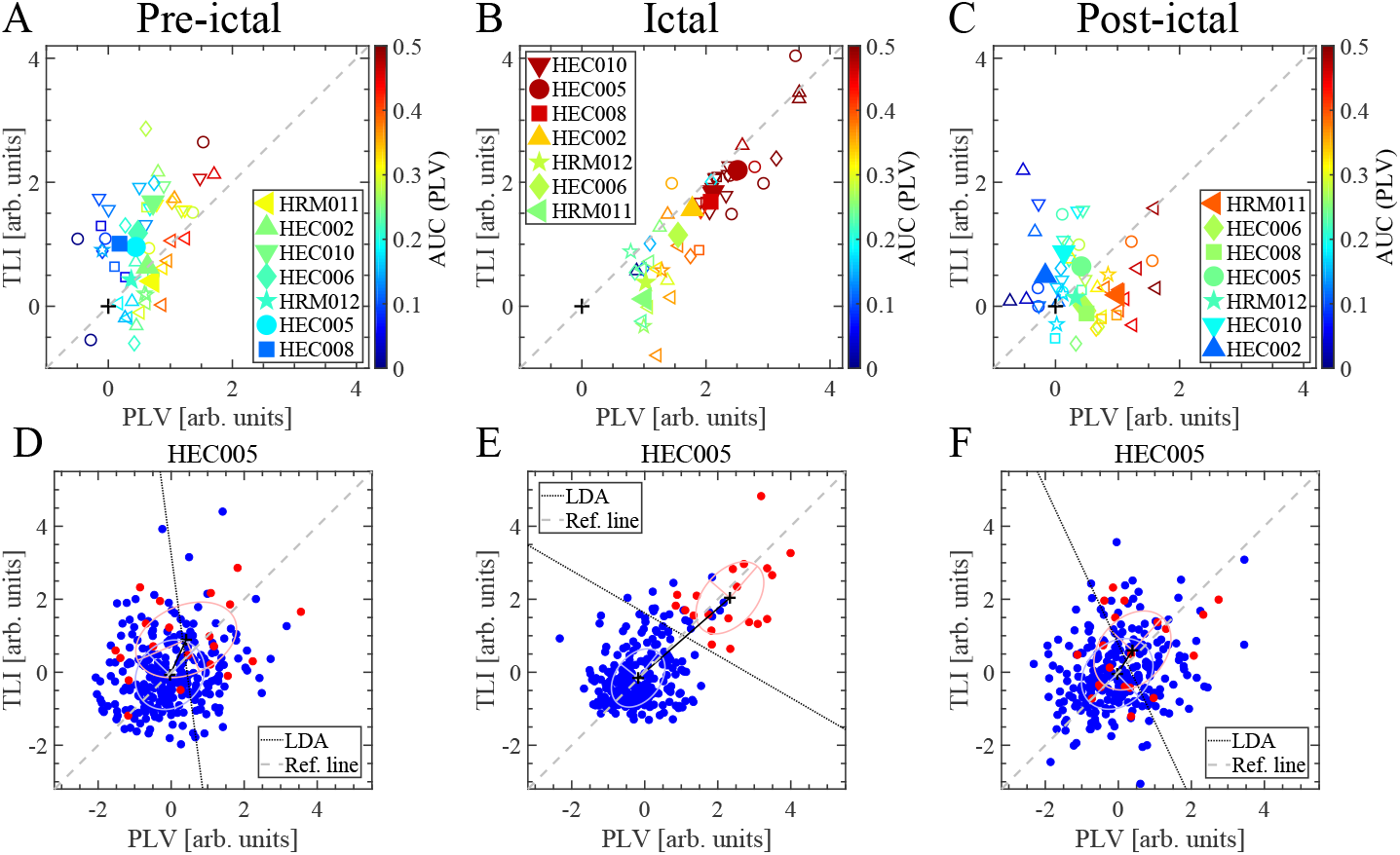
Bivariate analysis implemented for pre-ictal, ictal and post-ictal time intervals. (A, B, C) Euclidean distance between the centroids of the covariance error ellipses. The black cross in the origin and the colored markers represents the centroids of the non SOZ and SOZ covariance error ellipses, respectively. Filled markers were computed including all the seizures of each patient at once. On the other hand, the empty markers were computed for a given patient taking each seizure individually. (D, E, F) Seizure onset zone (SOZ, red filled circles) and non SOZ (blue filled circles) samples of Z-scored mean values in the PAC vs. harmonicity plane (PLV, TLI) including all the seizures of patient HEC005. Eigenvectors are shown together with the covariance error ellipses. The doted black line correspond to the Fisher’s discriminant resulting from the linear discriminant analysis (LDA). These graphs were computed for the frequency band combination (modulating LF vs. modulated HF) that maximizes the PLV-based SOZ classification power rated by the AUC in each patient (see Table A.2). Symbols and abbreviations: LF, low frequency; HF, high frequency; PAC, phase-amplitude coupling; PLV, phase locking value; TLI, time locked index; ROC, receiver operating characteristic; AUC, area under the ROC curve.

### 3.9. Intra-patient bivariate analysis

The results shown in Figures 14 and 15 were obtained by applying the bivariate analysis described in Section 2.10 computing in each patient a single frequency band combination (modulating LF vs. modulated HF) that maximizes the PLV-based iEEG SOZ classification power rated by the AUC in each patient (see Table A.2). In addition, the bivariate analysis described in Section 2.10 was also implemented in each patient for all the suitable combinations between the modulating LF (High Delta to High Beta) and modulated HF (Alpha to High HFO) frequency bands (see the inclusion criteria described in Section 2.9 and the frequency bands in the abscissa of Figures A.5, A.6 and A.7). Figure 16 shows the Euclidean distance between the centroids of the covariance error ellipses in the PAC vs. harmonicity plane (PLV, TLI) computed from the ictal time interval, including all the seizures in each patient and taking the frequency band pair as a parameter. In Figure 16, the black cross in the origin and the colored markers represents the centroids of the non SOZ and SOZ covariance error ellipses, respectively. Of note, the increase of the Euclidean distance between the centroids of the covariance error ellipses along the diagonal in the PAC vs. harmonicity plane (PLV, TLI), which is associated to the presence of harmonic ictal PAC, is present in both the inter-patient analysis (see Figures 14A and A.9A) and the intra-patient analysis (see the case HEC005 in Figure 16A). On the other hand, the behavior associated to the harmonic PAC is absent in the case HRM011 shown in Figure 16A, suggesting the presence of non harmonic ictal PAC in this patient. Moreover, it was found that the patterns shown in Figure 16 for the intra-patient analysis were preserved in the case of computing the seizures taken individually (data not shown).

**Figure 16:**
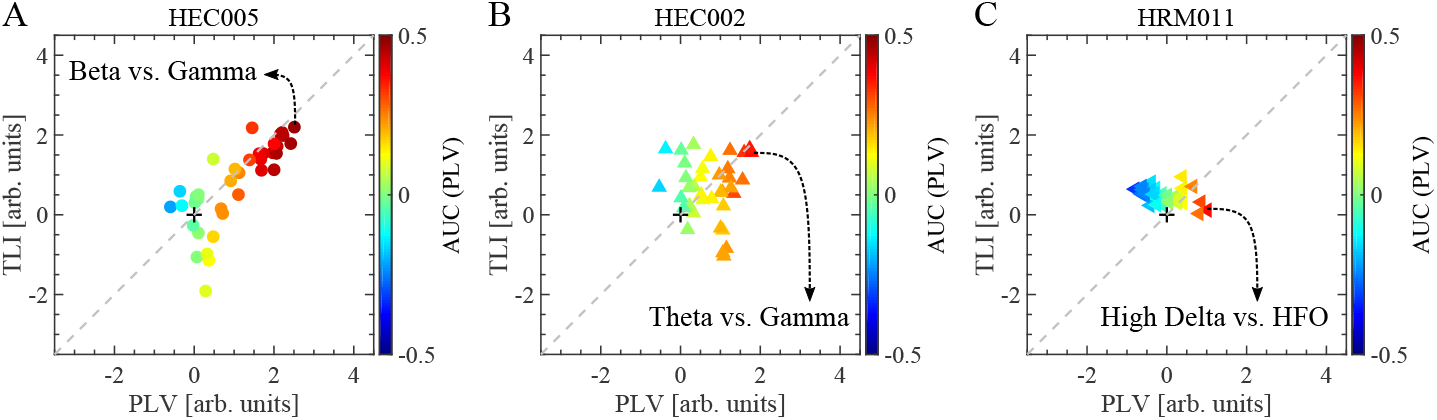
Intra-patient bivariate analysis implemented for the ictal time interval. The graphs show the Euclidean distance between the centroids of the covariance error ellipses in the PAC vs. harmonicity plane (PLV, TLI) computed from the ictal time interval, including all the seizures in each patient and taking the frequency band pair as a parameter (see the frequency bands indicated in the abscissa of Figures A.5C, A.6C and A.7C, and the inclusion criteria described in Section 2.9). The black cross in the origin and the colored markers represents the centroids of the non SOZ and SOZ covariance error ellipses, respectively. In each patient, it is indicated the frequency band combination (modulating LF vs. modulated HF) that maximizes the PLV-based SOZ classification power rated by the AUC (see Table A.2). Symbols and abbreviations: LF, low frequency; HF, high frequency; SOZ, seizure onset zone; PAC, phase-amplitude coupling; PLV, phase locking value; TLI, time locked index; ROC, receiver operating characteristic; AUC, area under the ROC curve.

## 4. DISCUSSION

We have demonstrated that, in the context of epileptic seizures, traditional algorithms aimed to assess PAC (e.g. PLV, KLMI) are confounded by harmonically related spectral components associated to non sinusoidal ictal activity, reporting significant PAC levels in absence of independent frequency bands (i.e. harmonic PAC). This is due to the fact that coupled oscillatory dynamics with independent frequencies (i.e. non harmonic PAC) and non sinusoidal ictal activity (i.e. harmonic PAC) produce similar signatures in the Fourier spectrum that are hardly distinguishable by using band-pass linear filtering.

### 4.1. Ictal harmonic PAC as a confounding factor

Recent studies have shown the relevance of PAC to detect epileptic seizures Ibrahim et al. (2014); Edakawa et al. (2016); Frauscher et al. (2015); Amiri et al. (2016) and it has been also hypothesized that ictal PAC, compared to interictal PAC, could provide more direct information to delineate the brain regions responsible for generation of habitual seizures Ibrahim et al. (2014); Motoi et al. (2018). More important, retrospective studies based on the postoperative surgical outcome of epilepsy surgery patients, have shown the clinical relevance of ictal PAC as a biomarker to improve the spatial delineation of resection boundaries associated to the ictal core from iEEG recordings with macroelectrodes Weiss et al. (2013, 2015); Schevon et al. (2012, 2019). Using invasive recordings with macro and microelectrodes in epilepsy patients, these studies demonstrated that repetitive transient increases in the amplitude of the high frequency band (80 Hz - 150 Hz), phase-locked to the slow ictal rhythm (1 Hz - 25 Hz), the so called phase-locked high gamma (PLHG), correlated with strong multi-unit firing bursts synchronized across the core territory of the seizure Weiss et al. (2013). This evidence indicates that the (non harmonic) PAC between the phase of low frequency modulating oscillations (High-Delta to Beta) and the amplitude of a modulated component pertaining to the High-Gamma and higher frequency bands, can be used as a surrogate of multi-unit firing bursts associated to the paroxysmal depolarizing shifts (PDSs) developed within the ictal core Weiss et al. (2013, 2015). The traditional PAC measures (e.g. PLV, KLMI) and, in principle, also the PLHG metric are susceptible to the harmonic PAC as a confounding factor since slow non sinusoidal rhythms (e.g. in the Delta frequency band) can produce prominent harmonically related spectral components, which extend inside the Gamma band (see Figures 4, 8 and 9). On the other hand, a recent work using non ictal invasive recordings has shown that coupled oscillations (i.e. non harmonic PAC) and repetitive sharp waveforms (i.e. harmonic PAC) emerging in the human cortex during an episodic memory task both produce PAC, however, they reflect two distinct neural mechanisms that are anatomically segregated in the human brain Vaz et al. (2017). As a consequence, the quantitative characterization of the harmonicity proposed in this work to effectively distinguish harmonic PAC from non harmonic PAC is relevant for the proper interpretation of the underlying ictal and physiological neural mechanisms.

### 4.2. Implications for improved therapy

The proposed harmonicity analysis is of valuable clinical utility since the harmonic PAC is a confounding factor which may lead to wrong conclusions about the underlying ictal mechanisms. That is, the paroxysmal depolarizing shifts (PDSs) and the restrained depolarizing shifts (RDSs) mechanisms of seizure propagation J.Trevelyan and A.Schevon (2013); Schevon et al. (2019), have both been associated to slow synaptic rhythms which are detectable with clinical macroelectrodes in the form of low frequency oscillatory LFP constituted by, in general, non sinusoidal waveform shapes (i.e. harmonic PAC). As a consequence, neither the low frequency LFP rhythms nor harmonic PAC patterns can be used to reliably differentiate the ictal core from the surrounding non fully recruited neural tissue (penumbra). Importantly, the hypersynchronized rhythmic burst firing coupled to the phase of slow synaptic rhythm, which is present only during the PDSs, produces a non harmonic PAC pattern with the modulated component pertaining to the High-Gamma and higher frequency bands. This non harmonic PAC pattern is specific of the fully recruited cortical areas (ictal core), constituting a clinically relevant biomarker that could be used to assist the spatial delineation of resection boundaries from iEEG recordings with macroelectrodes. The TLI metric made it possible to quantitatively differentiate non harmonic PAC from harmonic PAC patterns which coexist during the epileptiform activity in LFP recorded with invasive clinical macroelectrodes (see Figures 4, 8 and 9). Thus, the harmonicity analysis via the TLI provides valuable information that could be used to assess the extent of different ictal territories and the associated mechanisms of seizure propagation. A schematic representation of the PAC patterns associated to the PDSs and RDSs mechanisms discussed above is given in Figure 17.

**Figure 17:**
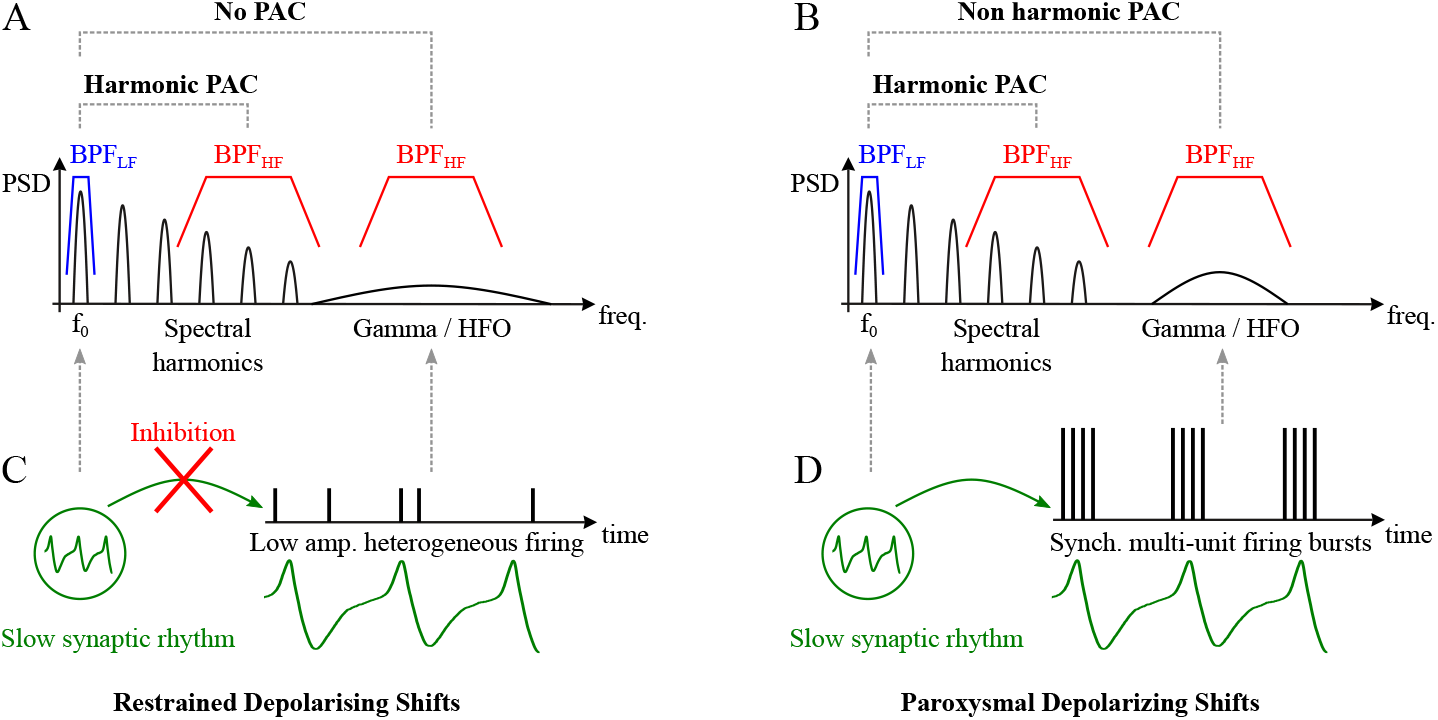
Schematic representation of the neural mechanisms and the associated CFC patterns. (A, B) Fourier spectrum and emergent harmonic and non harmonic PAC patterns observed in the LFP, associated to the mechanisms of seizure propagation. (C, D) Non sinusoidal slow synaptic rhythm (green curve) together with the spike patterns (vertical lines) associated to the mechanisms of seizure propagation termed as restrained depolarising shifts and paroxysmal depolarising shifts. Symbols and abbreviations: CFC, cross frequency coupling; PAC, phase-amplitude coupling; LFP, local field potential; HFO, high frequency oscillations; PSD, power spectral density; *f*_0_, fundamental frequency of the non sinusoidal oscillation; *BP F_LF_*, band pass filter for the low frequency band; *BP F_HF_*, band pass filter for the high frequency band.

### 4.3. The TLI as a complementary tool to assist the clinical analysis of iEEG recordings

It was found that in both inter-patient analysis (Figures 13, 14, 15 and A.9) and intra-patient analysis (Figure 16), the harmonicity as quantified by the TLI metric correlates with the performance of the ictal PAC to reproduce the classification of the iEEG SOZ given preoperatively by the epileptologists. Besides, all the studied patients undergoing resective epilepsy surgery (all but patient HEC010, see Table 1) present a good seizure outcome (i.e. Engel class I or II Weiss et al. (2015)). Taken together, these results suggest that the higher the harmonicity of the seizure activity, the better is the classification performance of the PAC biomarker to effectively identify the channels involved in the SOZ. As a conclusion, this evidence indicates that the analysis of how the bipolar iEEG channels are clustered in the PAC vs. harmonicity plane could be used to assist the visual-range analysis of the iEEG recordings performed by the epileptologists to define the SOZ (approx. 1 Hz - 30 Hz Schevon et al. (2019)). Besides, the temporal evolution of the PAC and harmonicity metrics is informative about the propagation of the seizure across the iEEG channels (see Figures 10 and 11). The latter is also of clinical relevance since, traditionally, the ictal core has been presumed to be identifiable from visualized initial ictal EEG changes (i.e. iEEG SOZ) Weiss et al. (2015).

### 4.4. Limitations and methodological considerations

This is a retrospective observational study, which should be considered in the interpretation of our results. Besides, although a variety of waveform shapes and PAC patterns were identified in the analyzed iEEG recordings, prospective and retrospective studies with a larger cohort of patients is warranted to effectively relate the ictal PAC and harmonicity patterns to the epileptogenic zones, and to the visual analysis performed by the epileptologists to define the SOZ. We emphasize that inter-area PAC and harmonicity analyses are also required to tackle the inverse problem associated to infer the underlying multidimensional neural dynamics using spatially sparse, one dimensional recordings.

## 5. CONCLUSION

We presented substantial evidence demonstrating, for the first time, the co-existence of harmonic PAC and non harmonic PAC patters during the seizure activity recorded from the iEEG SOZ of patients with pharmacoresistant focal epilepsy. This is relevant since traditional algorithms aimed to assess PAC are unable to distinguish between harmonic and non harmonic PAC patters observed during the seizure activity, which may lead to wrong conclusions about the underlying ictal mechanisms. Importantly, we found that a better putative SOZ classification based on the PAC biomarker, compared to the classification given preoperatively by the epileptologists, is accompanied by an increase in the harmonicity of the PAC pattern. Thus, the proposed PAC vs. harmonicity analysis is capable to provide clinically useful insights that may improve the effective delineation of the fully recruited cortical territories from invasive EEG recordings with macroelectrodes. As a conclusion, our results suggest that the spectral harmonicity analysis, using the TLI or any other measure, should be considered as a complementary tool to assist to the visual analysis of the iEEG recordings and to support the proper interpretation of ictal mechanisms associated to biomarkers based on cross frequency couplings.

## ACKNOWLEDGEMENTS

This work was partially supported by UNCuyo-SeCTyP 2019 (80020180100576UN, 80020180100653UN) Argentina.

## Appendix A. Supplementary methods

### Appendix A.1. Definition of the frequency bands considered in this study

**Table A.1:** Ranges for the frequency bands considered in this study.

| Band<br>name | Frequency<br>range [Hz] |
| --- | --- |
| Low Delta | 0.1 - 1 |
| High Delta | 1 - 4 |
| Delta | 0.1 - 4 |
| Theta | 4 - 8 |
| Alpha | 8 - 13 |
| Low Beta | 13 - 18 |
| High Beta | 18 - 30 |
| Beta | 13 - 30 |
| Low Gamma | 30 - 60 |
| High Gamma | 60 - 100 |
| Gamma | 30 - 100 |
| Low HFO | 100 - 250 |
| High HFO | 250 - 500 |
| HFO | 100 - 500 |
| Abbreviations: HFO, high frequency oscillations. |  |

### Appendix A.2. Two groups analysis

A two group statistical analysis was performed for the PAC (PLV, KLMI) and the harmonicity (TLI) time series. These time series were computed as described in Section 2.7. Importantly, the temporal correlation between the PLV and TLI time series was also included in the two group analysis, since the increase of the PAC intensity (PLV, KLMI) simultaneously with an increase of the TLI metric points out the presence of ‘harmonic PAC’ elicited by harmonic spectral components in the modulating LF and modulated HF frequency bands. The temporal correlation between the PAC metrics (PLV, KLMI) and the harmonicity metric (TLI) was assessed by computing the Spearman’s rank correlation coefficient between the corresponding time series (e.g. temporal correlation between the PLV and TLI time series). The two groups analysis was implemented for each patient as follows. In each single seizure, the PLV time series (one per bipolar iEEG channel) were segregated in two groups (SOZ and non SOZ) following the iEEG SOZ classification given preoperatively by the neurologist. Then, the SOZ and non SOZ groups were merged across all the seizures. As a result, we obtained two sample distributions of PLV values labeled as SOZ and non SOZ. A similar procedure was implemented for the TLI time series and for the temporal correlation between the PLV and TLI time series. In Figures 13A, A.1, A.2 and A.3 the sample distributions of PLV, TLI and PLV vs. TLI temporal correlation values pertaining to SOZ iEEG channels and non SOZ iEEG channels are represented as red and blue violin plots, respectively.

**Figure A.1:**
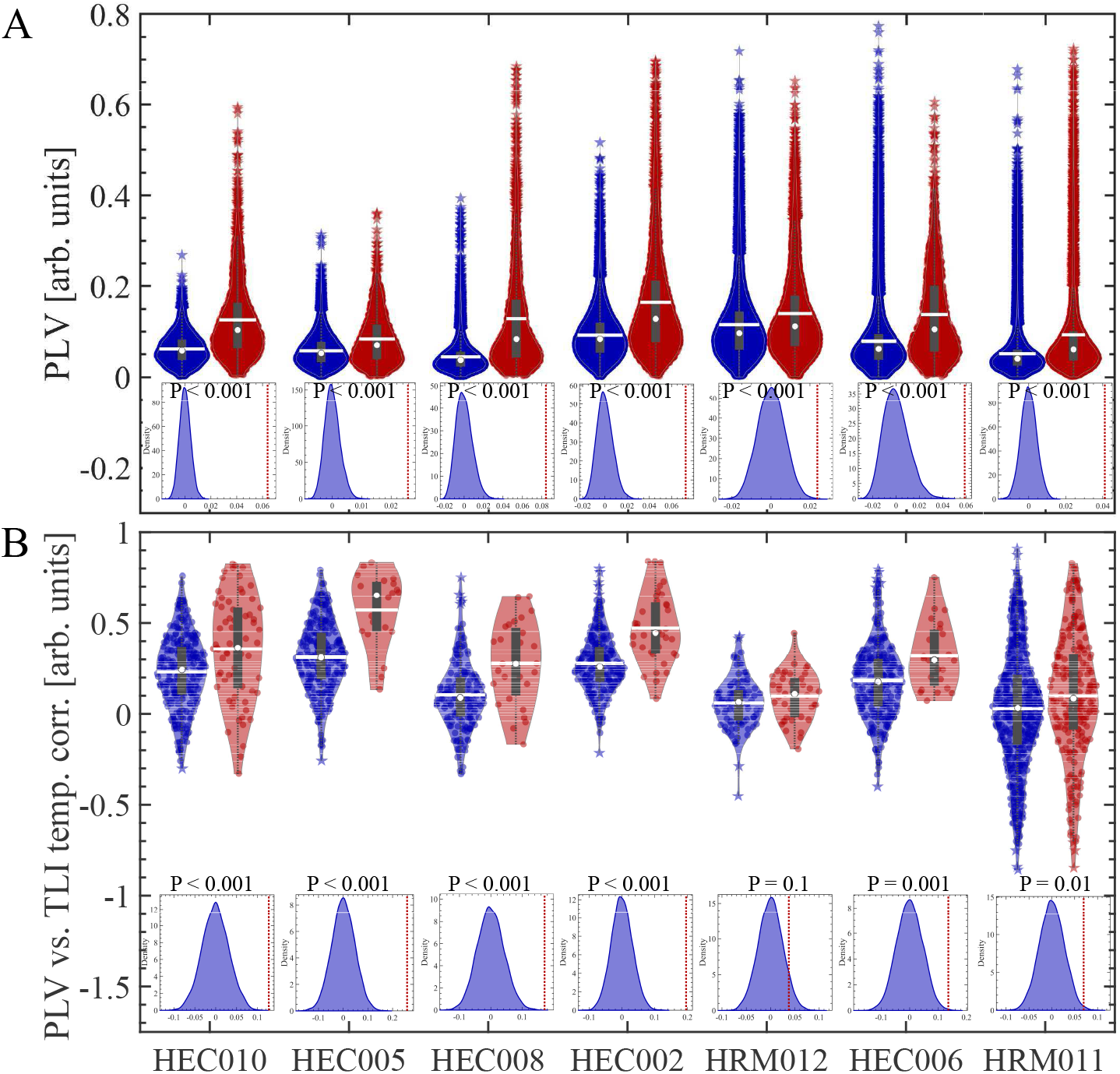
Two group analysis implemented for the ictal time interval (i.e. spanning the entire duration of each seizure). (A) Red and blue violin plots correspond to the SOZ and non SOZ sample distributions of PLV values, respectively. (B) Red and blue violin plots correspond to the SOZ and non SOZ sample distributions of PLV vs. TLI temporal correlation values, respectively. The PLV and TLI metrics were used to quantify PAC and harmonicity, respectively. In the violin plots, center gray boxes represent the 25th and 75th percentiles, whiskers (doted gray lines) extend to the most extreme data points not considered outliers (1.5 times the interquartile range), star markers represent outliers. The center white circle and white line indicate the median and mean, respectively. The histograms correspond to the sampling distribution of the difference between the means of the SOZ and non SOZ distributions. The histograms were computed via random sampling without replacement (1 10^4^ permutations). The vertical doted line (red) in the histograms indicates the actual mean difference between the SOZ and non SOZ distributions shown in the corresponding violin plots. These graphs were computed for the frequency band combination (modulating LF vs. modulated HF) that maximizes the PLV-based SOZ classification power rated by the AUC in each patient (see Table A.2). Symbols and abbreviations: LF, low frequency; HF, high frequency; SOZ, seizure onset zone; PAC, phase-amplitude coupling; PLV, phase locking value; TLI, time locked index; ROC, receiver operating characteristic; AUC, area under the ROC curve.

**Figure A.2:**
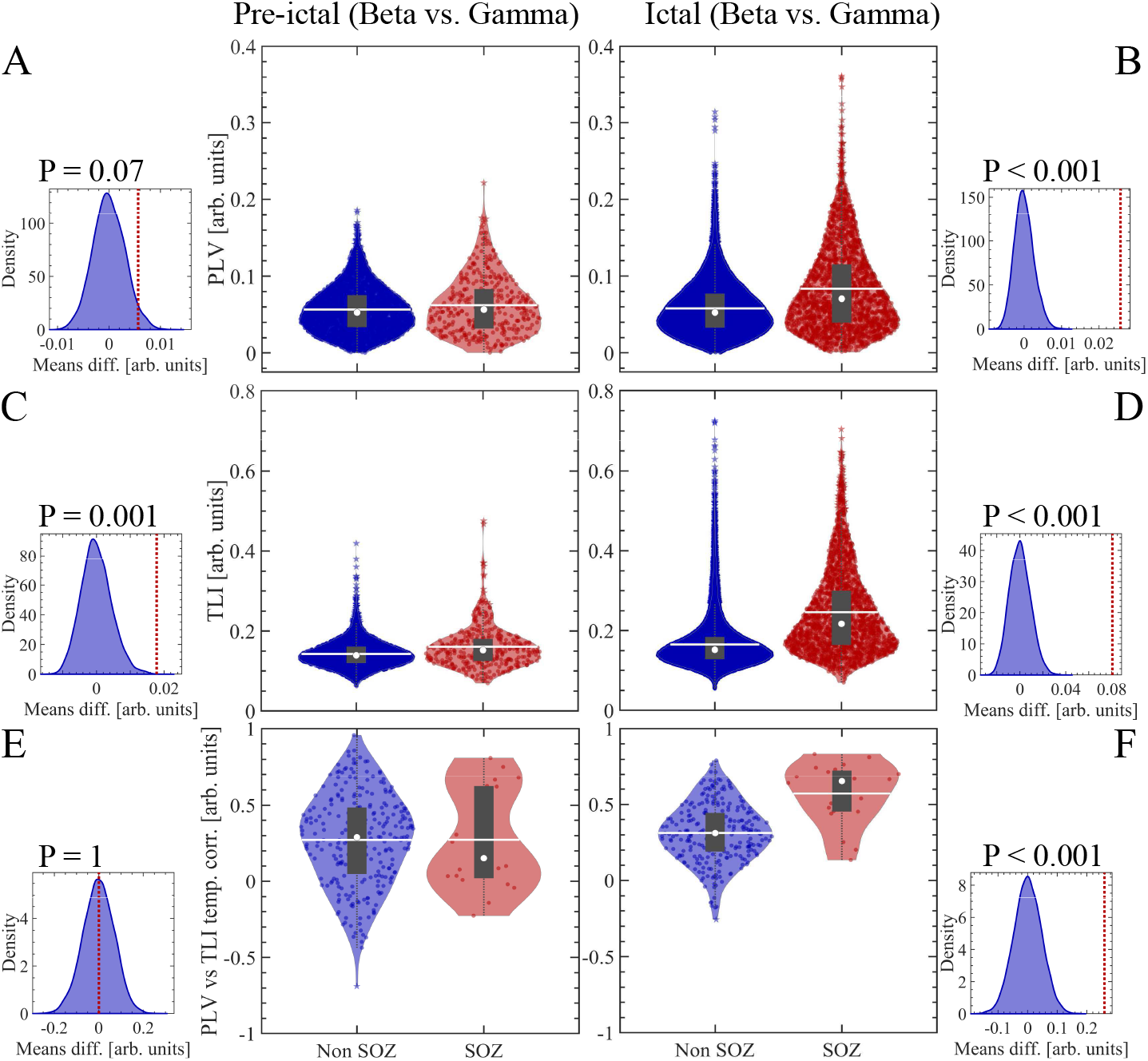
Two group analysis implemented for pre-ictal (panels A, C, E) and ictal (panels B, D, F) time intervals in patient HEC005. (A, B) Red and blue violin plots correspond to the SOZ and non SOZ sample distributions of PLV values, respectively. (C, D) Red and blue violin plots correspond to the SOZ and non SOZ sample distributions of TLI values, respectively. (E, F) Red and blue violin plots correspond to the SOZ and non SOZ sample distributions of PLV vs. TLI temporal correlation values, respectively. The PLV and TLI metrics were used to quantify PAC and harmonicity, respectively. In the violin plots, center gray boxes represent the 25th and 75th percentiles, whiskers (doted gray lines) extend to the most extreme data points not considered outliers (1.5 times the interquartile range), star markers represent outliers. The center white circle and white line indicate the median and mean, respectively. The histograms correspond to the sampling distribution of the difference between the means of the SOZ and non SOZ distributions. The histograms were computed via random sampling without replacement (1 10^4^ permutations). The vertical doted line (red) in the histograms indicates the actual mean difference between the SOZ and non SOZ distributions shown in the corresponding violin plots. These graphs were computed for the frequency band combination (modulating LF vs. modulated HF) that maximizes the PLV-based SOZ classification power rated by the AUC for the patient HEC005 (see Table A.2). Symbols and abbreviations: LF, low frequency; HF, high frequency; SOZ, seizure onset zone; PAC, phase-amplitude coupling; PLV, phase locking value; TLI, time locked index; ROC, receiver operating characteristic; AUC, area under the ROC curve.

**Figure A.3:**
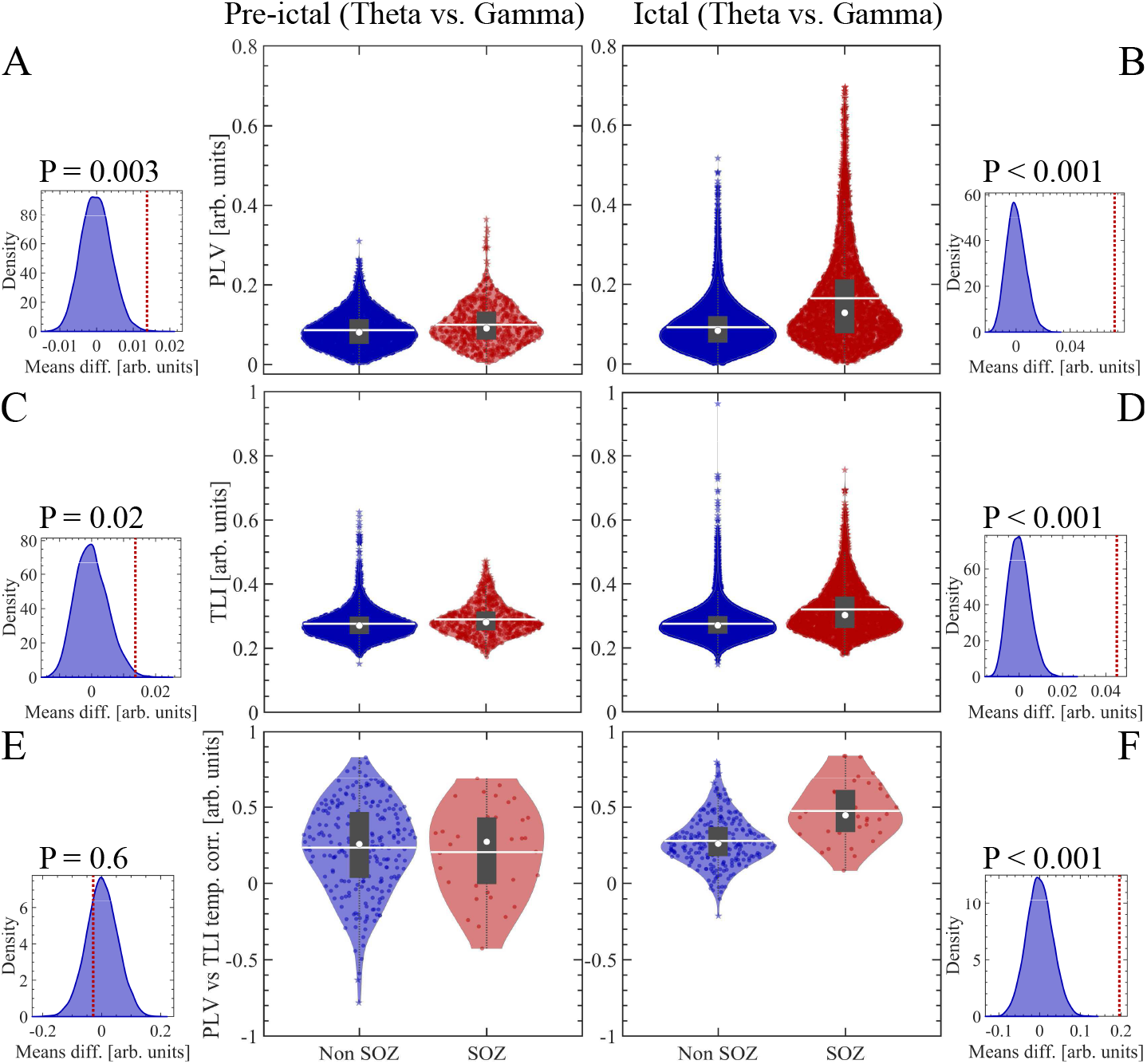
Two group analysis implemented for pre-ictal (panels A, C, E) and ictal (panels B, D, F) time intervals in patient HEC002. (A, B) Red and blue violin plots correspond to the SOZ and non SOZ sample distributions of PLV values, respectively. (C, D) Red and blue violin plots correspond to the SOZ and non SOZ sample distributions of TLI values, respectively. (E, F) Red and blue violin plots correspond to the SOZ and non SOZ sample distributions of PLV vs. TLI temporal correlation values, respectively. The PLV and TLI metrics were used to quantify PAC and harmonicity, respectively. In the violin plots, center gray boxes represent the 25th and 75th percentiles, whiskers (doted gray lines) extend to the most extreme data points not considered outliers (1.5 times the interquartile range), star markers represent outliers. The center white circle and white line indicate the median and mean, respectively. The histograms correspond to the sampling distribution of the difference between the means of the SOZ and non SOZ distributions. The histograms were computed via random sampling without replacement (1 10^4^ permutations). The vertical doted line (red) in the histograms indicates the actual mean difference between the SOZ and non SOZ distributions shown in the corresponding violin plots. These graphs were computed for the frequency band combination (modulating LF vs. modulated HF) that maximizes the PLV-based iEEG SOZ classification power rated by the AUC for the patient HEC002 (see Table A.2). Symbols and abbreviations: LF, low frequency; HF, high frequency; SOZ, seizure onset zone; PAC, phase-amplitude coupling; PLV, phase locking value; TLI, time locked index; ROC, receiver operating characteristic; AUC, area under the ROC curve.

### Appendix A.3. Paired analysis

The paired analysis for the PAC and harmonicity metrics was implemented for each patient as follows. Pre-ictal and ictal time series for each bipolar iEEG channel were constructed for the PLV metric as described in Section 2.7. To this purpose, the PLV metric was computed for the frequency band combination (modulating LF vs. modulated HF) that maximizes the PLV-based iEEG SOZ classification power rated by the AUC in each patient (see Table A.2). Then, we assign to each bipolar iEEG channel a pair of PLV mean values computed form the pre-ictal and ictal PLV time series. After that, the pre-ictal and ictal pairs of PLV mean values were segregated in the SOZ and non SOZ groups which were merged across all the seizures following a procedure similar to that described in Appendix A.2. This series of steps was also used to implement the paired analysis for the TLI metric. The statistical significance of the paired mean values (PLV or TLI) corresponding to the pre-ictal and ictal time intervals, was assessed using the Wilcoxon signed rank test including the Bonferroni correction for multiple comparisons.

**Figure A.4:**
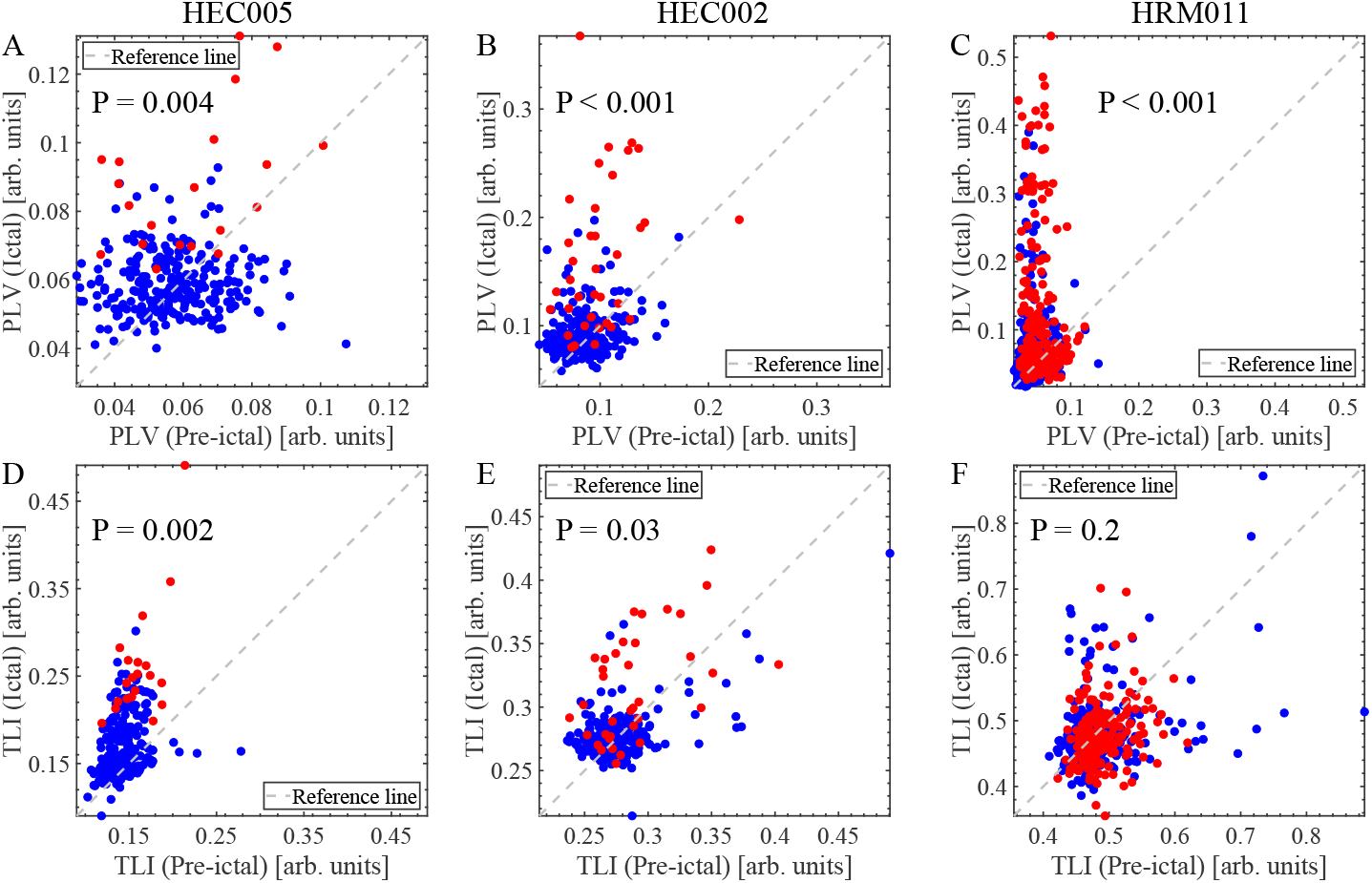
Scatter plots for the paired analysis. (A, B, C) Scatter plots (Ictal vs. Pre-ictal) of PLV mean values including all bipolar channels in each patient. (D, E, F) Scatter plots (Ictal vs. Pre-ictal) of TLI mean values including all bipolar channels in each patient. Red and blue markers represent mean values (PLV, TLI) computed from bipolar channels pertaining to SOZ and non SOZ, respectively. These graphs were computed for the frequency band combination (modulating LF vs. modulated HF) that maximizes the PLV-based SOZ classification power rated by the AUC in each patient (see Table A.2). The Bonferroni-adjusted P values reported in the graphs were obtained using the Wilcoxon signed rank test for the bipolar channels pertaining to the SOZ (red markers). Symbols and abbreviations: LF, low frequency; HF, high frequency; SOZ, seizure onset zone; PAC, phase-amplitude coupling; PLV, phase locking value; TLI, time locked index; ROC, receiver operating characteristic; AUC, area under the ROC curve.

### Appendix A.4. Receiver operating characteristic analysis

In the following we present the results obtained with the receiver operating characteristic (ROC) analysis implemented as it was described in Section 2.9. Figures A.5, A.6 and A.7 show the results of the ROC analysis computed from PLV and TLI time series spanning the entire duration of each seizure (i.e. ictal time series) for the patients HEC005, HEC002 and HRM011, respectively. For the patient HEC005, Figure A.5A shows the mean values of the ictal time series of the PLV metric from which the Beta vs. Gamma ROC curve was computed by applying a moving threshold on the PLV intensity (see the ROC curve in Figure A.5B). Note that in computing the ROC curves we have one mean value per seizure and bipolar iEEG channel (7 seizures for the patient HEC005, see Table 1). Figure A.5C shows the AUC for PLV and TLI metrics as functions of the frequency band combination (modulating LF vs. modulated HF). Beta vs. Gamma is the pair of modulating LF vs. modulated HF frequency band combination that maximizes the iEEG SOZ classification power of PLV metric rated by the AUC. This frequency band combination is highlighted by the arrow in Figure A.5C. In Figure A.5C, it is possible to distinguish local maxima with similar AUC values for the PLV metric occurring around the modulating LF frequency bands Theta, Alpha, Low Beta, Beta and High Beta. Moreover, the AUC values corresponding to the TLI metric closely reproduce this pattern across the frequency bands. Importantly, every time AUC for PLV metric is high, AUC for TLI metric is high as well. This behavior emerges from the presence of harmonic PAC and can be interpreted as follows. The harmonic spectral components produced by the quasiperiodic non sinusoidal ictal activity span several frequency bands (from Theta to Low HFO) giving rise to significant harmonic PAC values as measured by the PLV metric (see Figures 5, 4 and 8). Hence, an increase of harmonicity in the bipolar iEEG channels pertaining SOZ relative to the rest of the channels, produces a similar relative increase of the mean values of TLI and PLV intensity when computed in the frequency bands capturing the spectral harmonics, and thus, resulting in high AUC values (AUC ≈ 0.5) for those frequency bands. The AUC values shown in Figure A.5 for both, the PLV and TLI metrics are close to the chance level (AUC ≈ 0) in the case of the modulating High Delta band suggesting the absence of spectral harmonics associated to this frequency band, which is consistent with the spectrogram and spectrum presented in Figure 4. For the patient HRM011, Figure A.7A shows the mean values of the ictal time series of the PLV metric from which the High Delta vs. HFO ROC curve was computed by applying a moving threshold on the PLV intensity (see the ROC curve in Figure A.7B). Note that in computing the ROC curves we have one mean value per seizure and bipolar iEEG channel (9 seizures for the patient HRM011, see Table 1). Figure A.7C shows the AUC for PLV and TLI metrics as functions of the frequency band combination (modulating LF vs. modulated HF). High Delta vs. HFO is the pair of modulating LF vs. modulated HF frequency band combination that maximizes the iEEG SOZ classification power of PLV metric rated by the AUC. This frequency band combination is highlighted by the arrow in Figure A.7C. Note that the maximum AUC value for the PLV metric co-occur with the minimum AUC value for the TLI metric at High Delta vs. HFO (arrow in Figure A.7C). The latter supports the hypothesis discussed in Section 3.1 regarding the presence of non harmonic PAC in the ictal activity of this patient.

The frequency bands maximizing the iEEG SOZ classification rated by the AUC obtained from the ROC analysis of the power and PAC measures are listed for the seven patients in Table A.2. Figure A.8 shows the AUC values for PLV and TLI metrics computed for the frequency band combinations which maximizes the PLV-based iEEG SOZ classification (see Table A.2). Of note, the AUC values for PLV and TLI metrics shown in Figure A.8 are highly correlated suggesting that the performance of PAC for iEEG SOZ classification is closely related to the harmonicity of the ictal activity.

**Figure A.5:**
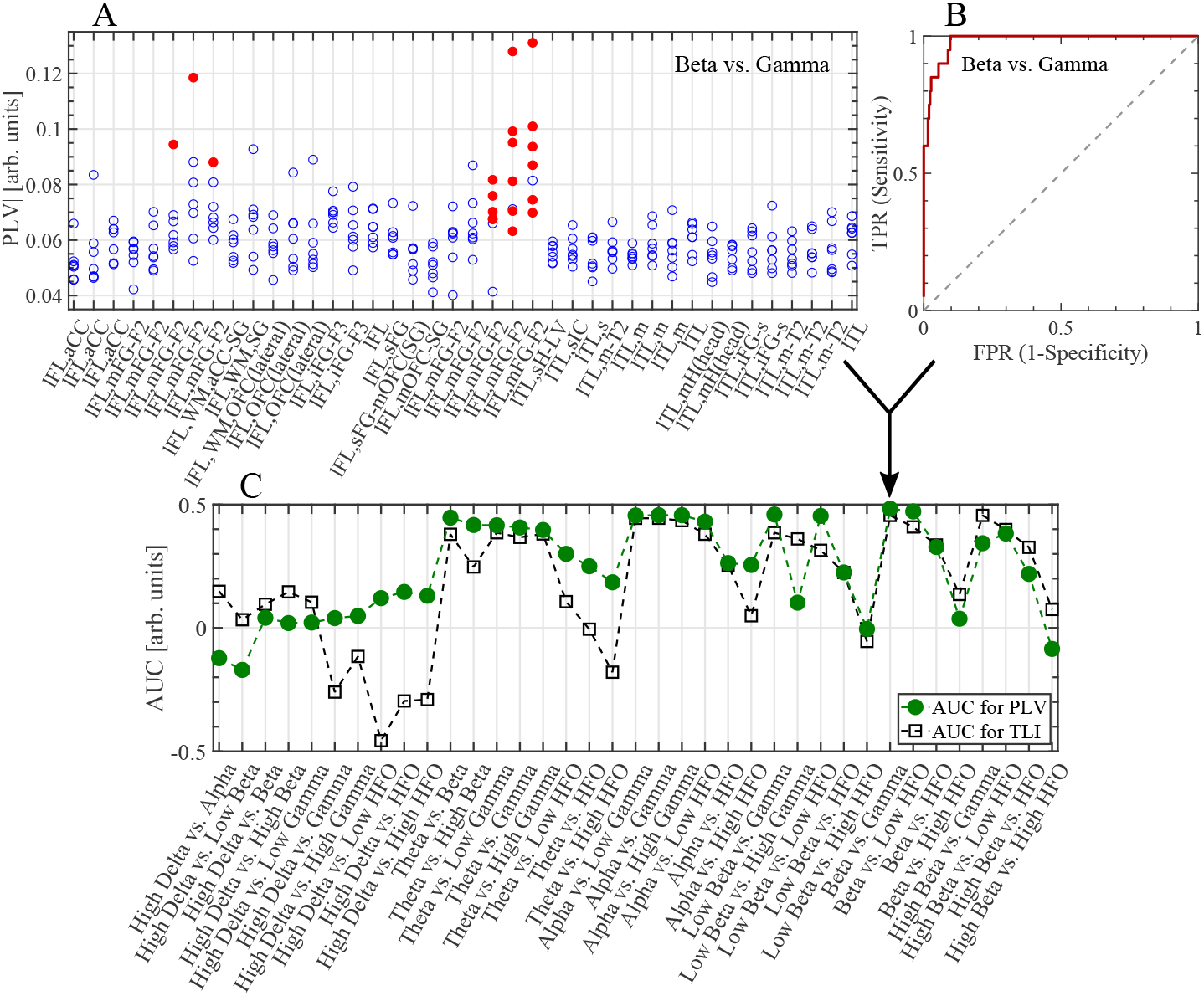
Receiver operating characteristic (ROC) analysis for patient HEC005. (A) Mean values of phase locking value (PLV) magnitude for quantification of phase-amplitude coupling (PAC) during the ictal time interval for the Beta vs. Gamma frequency band combination (modulating LF vs. modulated HF). Filled red and empty blue markers represent mean values of PLV magnitude computed from recordings pertaining to seizure onset zone (SOZ) and non SOZ, respectively. Note that the 7 points per bipolar channel correspond to the number of seizures processed for this patient (see Table 1). The labels of bipolar channels of patient HEC005 are described in Table A.3. (B) ROC curve for the Beta vs. Gamma frequency band combination, obtained by applying a moving threshold on the mean values of PLV magnitude (panel A). (C) Area under the ROC curve (AUC) as a function of the frequency band combinations for the PLV and time locked index (TLI) metrics. Note that Beta vs. Gamma (black arrow) is the frequency band combination that maximizes the AUC for PLV in this patient (AUC_PLV_=0.48, see Table A.2). Symbols and abbreviations: LF, low frequency; HF, high frequency; HFO, high frequency oscillations; TPR, true positive rate; FPR, false positive rate; AUC_PLV_, AUC for PAC computed using the PLV metric.

**Figure A.6:**
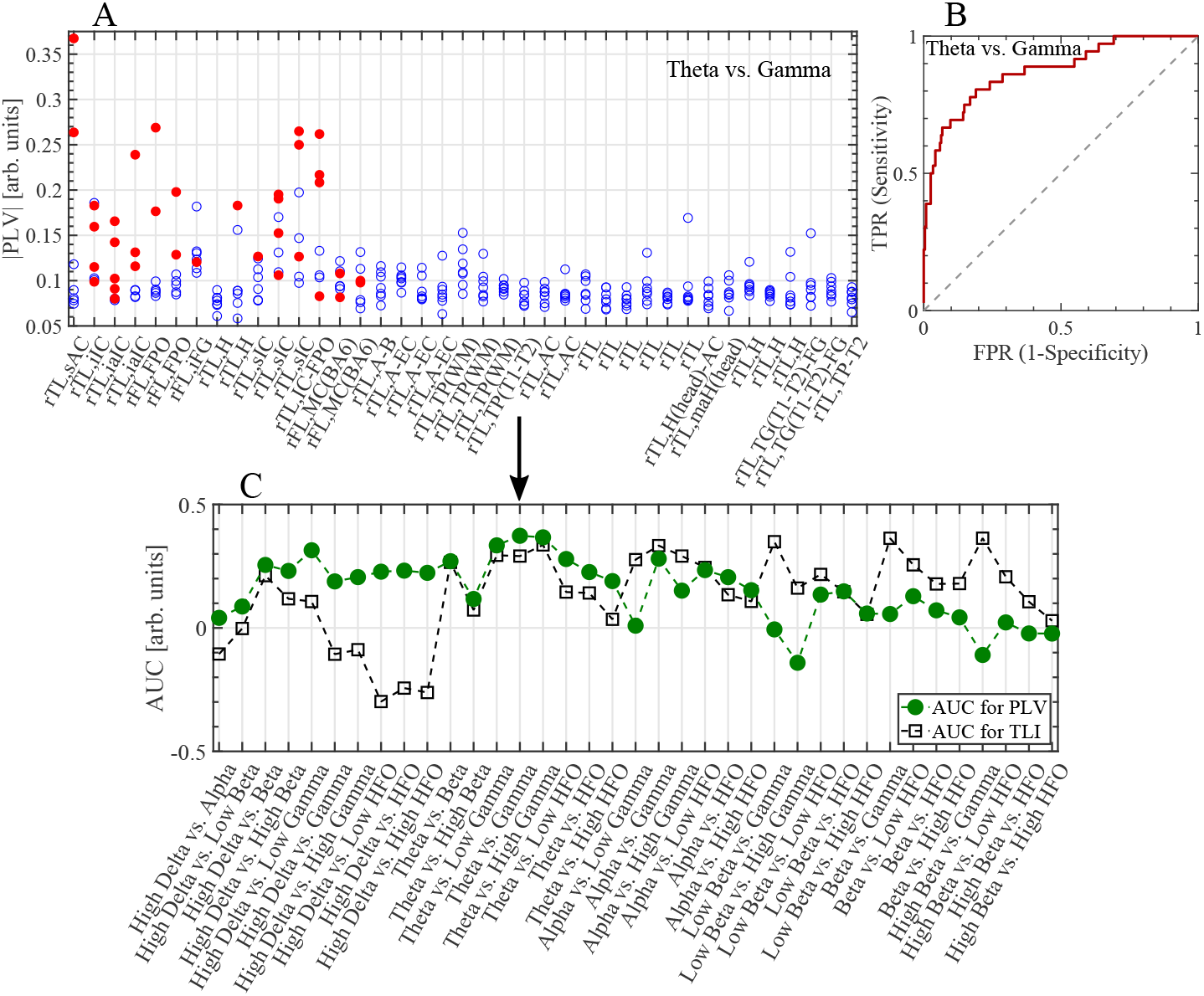
Receiver operating characteristic (ROC) analysis for patient HEC002. (A) Mean values of phase locking value (PLV) magnitude during the ictal time interval for the Theta vs. Gamma frequency band combination (modulating LF vs. modulated HF). Filled red and empty blue markers represent mean values of PLV magnitude computed from recordings pertaining to seizure onset zone (SOZ) and non SOZ, respectively. Note that the 7 points per bipolar channel correspond to the number of seizures processed for this patient (see Table 1). The labels of bipolar channels of patient HEC002 are described in Table A.4. (B) ROC curve for the Theta vs. Gamma frequency band combination, obtained by applying a moving threshold on the mean values of PLV magnitude (panel A). (C) Area under the ROC curve (AUC) as a function of the frequency band combinations for the PLV and time locked index (TLI) metrics. Note that Theta vs. Gamma (black arrow) is the frequency band combination that maximizes the AUC for PLV in this patient (AUC_PLV_=0.37, see Table A.2). Symbols and abbreviations: LF, low frequency; HF, high frequency; HFO, high frequency oscillations; TPR, true positive rate; FPR, false positive rate; AUC_PLV_, AUC for PAC computed using the PLV metric.

**Figure A.7:**
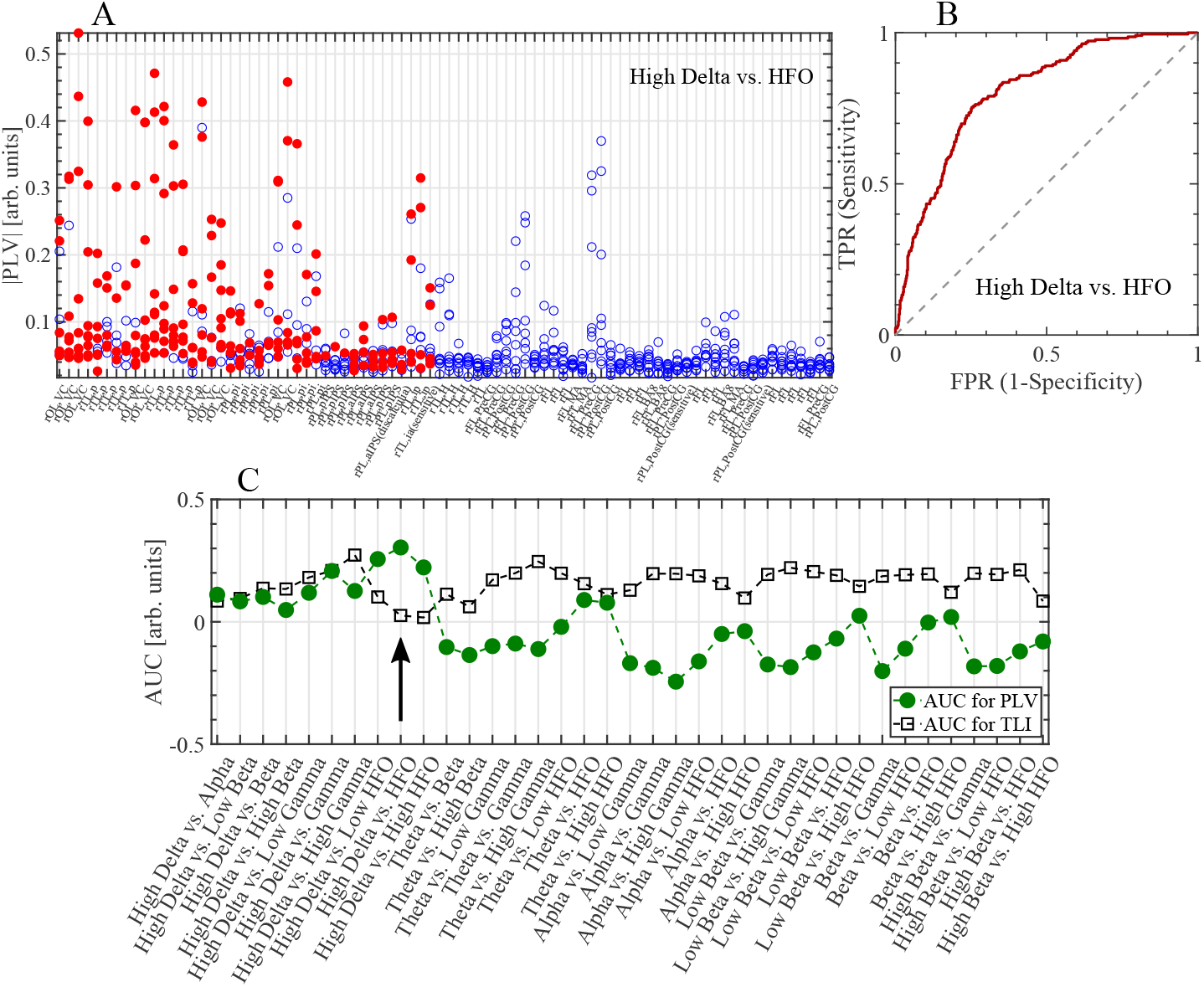
Receiver operating characteristic (ROC) analysis for patient HRM011. (A) Mean values of phase locking value (PLV) magnitude during the ictal time interval for the High Delta vs. HFO frequency band combination (modulating LF vs. modulated HF). Filled red and empty blue markers represent mean values of PLV magnitude computed from recordings pertaining to seizure onset zone (SOZ) and non SOZ, respectively. Note that the 9 points per bipolar channel correspond to the number of seizures processed for this patient (see Table 1). The labels of bipolar channels of patient HRM011 are described in Table A.5. (B) ROC curve for the High Delta vs. HFO frequency band combination, obtained by applying a moving threshold on the mean values of PLV magnitude (panel A). (C) Area under the ROC curve (AUC) as a function of the frequency band combinations for the PLV and time locked index (TLI) metrics. Note that High Delta vs. HFO (black arrow) is the frequency band combination that maximizes the AUC for PLV in this patient (AUC_PLV_=0.30, see Table A.2). Symbols and abbreviations: LF, low frequency; HF, high frequency; HFO, high frequency oscillations; TPR, true positive rate; FPR, false positive rate; AUC_PLV_, AUC for PAC computed using the PLV metric.

**Figure A.8:**
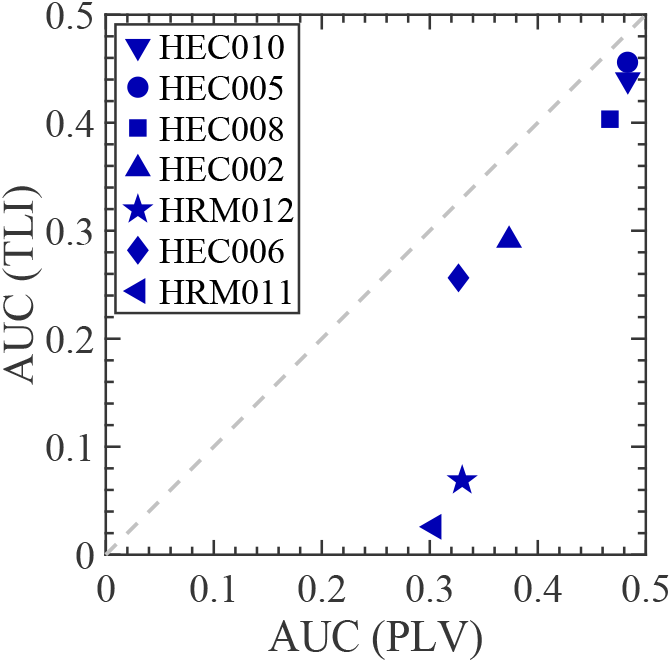
AUC values for PAC (PLV) and harmonicity (TLI) metrics computed for the frequency band combination (modulating LF vs. modulated HF) that maximizes the PLV-based SOZ classification (see Table A.2). Pearson’s correlation analysis confirmed a highly significant linear correlation between the AUC for PAC using PLV and AUC for harmonicity using TLI (r = 0.92, P = 0.003, t(5) = 5.2, Student’s t-test). Symbols and abbreviations: LF, low frequency; HF, high frequency; SOZ, seizure onset zone; PAC, phase-amplitude coupling; PLV, phase locking value; TLI, time locked index; ROC, receiver operating characteristic; AUC, area under the ROC curve.

**Table A.2:** Frequency bands maximizing the iEEG SOZ classification rated by the AUC obtained from the ROC analysis considering the ictal time interval.

| Patient | Frequency band<br>for Power ($AUC_{Power}$) | Frequency bands<br>for PAC ($AUC_{PLV}$) |
| --- | --- | --- |
| HEC010 | High Gamma (0.49) | Alpha vs. Low HFO (0.48) |
| HEC005 | Beta (0.45) | Beta vs. Gamma (0.48) |
| HEC008 | Low Delta (0.38) | Theta vs. HFO (0.47) |
| HEC002 | Beta (0.36) | Theta vs. Gamma (0.37) |
| HRM012 | HFO (0.29) | High Delta vs. Gamma (0.33) |
| HEC006 | High Beta (0.33) | Theta vs. Low HFO (0.33) |
| HRM011 | HFO (0.31) | High Delta vs. HFO (0.30) |

Abbreviations: iEEG SOZ, seizure onset zone identified from the intracerebral electroencephalography recordings;
PLV, phase locking value;
PAC, phase-amplitude couplings;
ROC, receiver operating characteristic;
AUC, area under the ROC curve;
$AUC_{Power}$ , AUC computed using the spectral power metric;
$AUC_{PLV}$ , AUC for PAC computed using the PLV metric;
HFO, high frequency oscillations.

### Appendix A.5. Bivariate analysis

Figures A.9B and A.9C show the SOZ (red filled circles) and non SOZ (blue filled circles) samples of Z-scored mean values in the PAC vs. harmonicity plane (KLMI, TLI) for the patients HEC005 and HRM011, which were computed for the ictal time interval and the frequency band combination (modulating LF vs. modulated HF) that maximizes the PLV-based iEEG SOZ classification power rated by the AUC in each patient (see Table A.2). Figure A.9A shows the Euclidean distance between the centroids of the covariance error ellipses in the PAC vs. harmonicity plane (KLMI, TLI) computed from the ictal time interval for the seven patients. In Figure A.9A, the black cross in the origin and the colored markers represents the centroids of the non SOZ and SOZ covariance error ellipses, respectively. It is worth noting that Figures A.9B and A.9C and the filled markers in Figure A.9A were computed including all the seizures of each patient at once, as described in Section 2.10. On the other hand, the empty markers in Figure A.9A were obtained by applying the same procedure over each seizure individually. That is, we obtained an Euclidean distance between the centroids of the covariance error ellipses in the PAC vs. harmonicity plane (KLMI, TLI) for each single seizure of a given patient. Importantly, Figure A.9 computed using the KLMI metric reproduces the behavior observed in the Figure 14 computed using the PLV metric, suggesting that this result is highly independent of the metric used to quantify PAC.

**Figure A.9:**
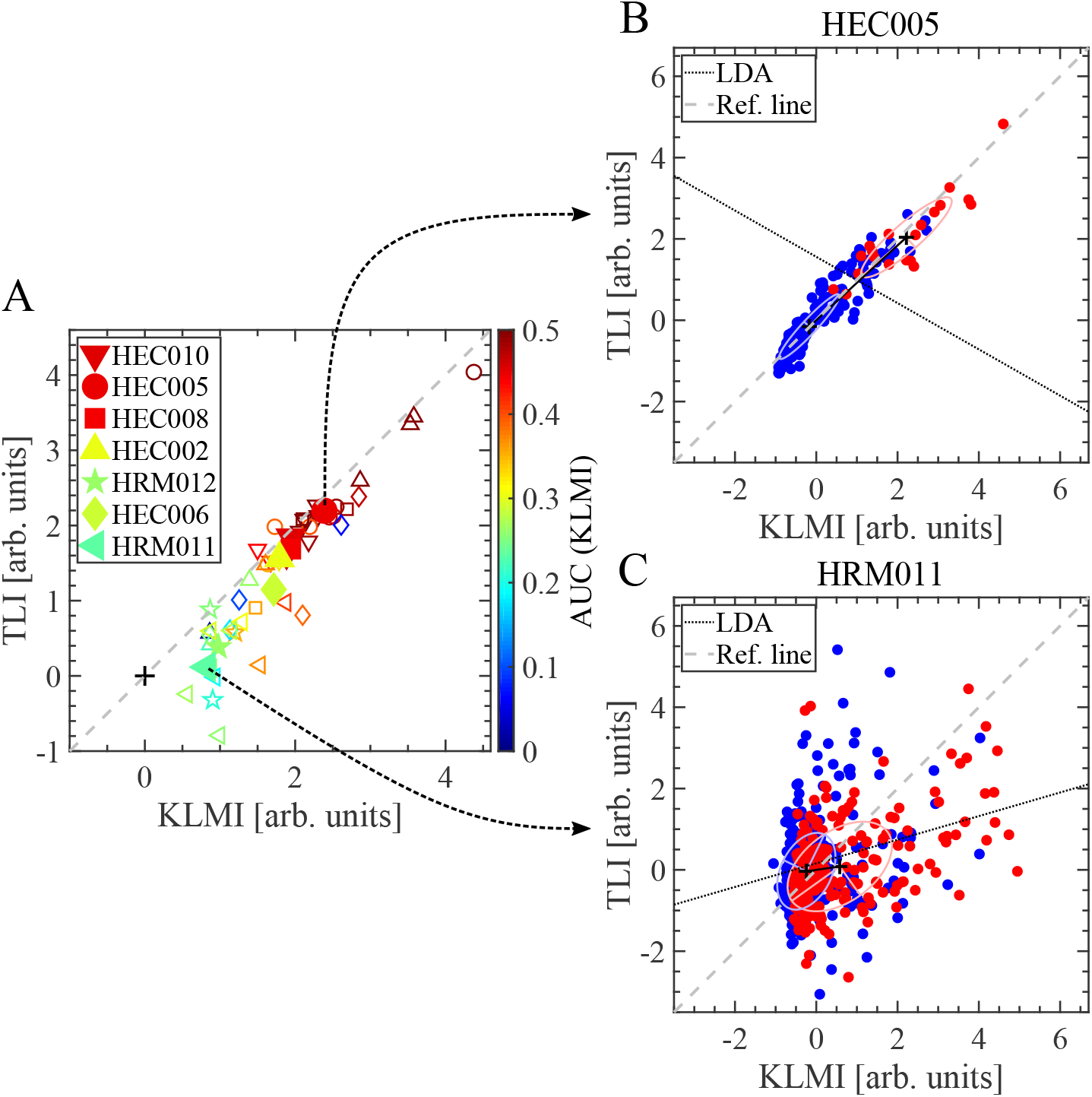
Bivariate analysis implemented for the ictal time interval (i.e. spanning the entire duration of each seizure). (A) Euclidean distance between the centroids of the covariance error ellipses. The black cross in the origin and the colored markers represents the centroids of the non SOZ and SOZ covariance error ellipses, respectively. Filled markers were computed including all the seizures of each patient at once. On the other hand, the empty markers were computed for a given patient taking each seizure individually. (B, C) Seizure onset zone (SOZ, red filled circles) and non SOZ (blue filled circles) samples of Z-scored mean values in the PAC vs. harmonicity plane (KLMI, TLI) including all the seizures of each patient. Eigenvectors are shown together with the covariance error ellipses. The doted black line correspond to the Fisher’s discriminant resulting from the linear discriminant analysis (LDA) and the dashed gray line is the reference line. These graphs were computed for the frequency band combination (modulating LF vs. modulated HF) that maximizes the PLV-based SOZ classification power rated by the AUC in each patient (see Table A.2). Symbols and abbreviations: LF, low frequency; HF, high frequency; PAC, phase-amplitude coupling; KLMI, modulation index based on the Kullback-Leibler distance; PLV, phase locking value; TLI, time locked index; ROC, receiver operating characteristic; AUC, area under the ROC curve.

### Appendix A.6. Labels describing the anatomical location of bipolar recordings

**Table A.3:** Labels describing the anatomical location of bipolar recordings from patient HEC005.

| Label | Description |
| --- | --- |
| lFL | left frontal lobe |
| lFL,aCC | left frontal lobe, anterior cingulate cortex |
| lFL,mFG-F2 | left frontal lobe, middle frontal gyrus - area F2 |
| lFL,WM,aCC-SG | left frontal lobe, white matter, anterior cingulate cortex - straight gyrus |
| lFL,WM,SG | left frontal lobe, white matter, straight gyrus |
| lFL,WM,OFC(lateral) | left frontal lobe, white matter, lateral orbitofrontal cortex |
| lFL,OFC(lateral) | left frontal lobe, lateral orbitofrontal cortex |
| lFL,iFG-F3 | left frontal lobe, inferior frontal gyrus - area F3 |
| lFL,sFG | left frontal lobe, superior frontal gyrus |
| lFL,sFG-mOFC(SG) | left frontal lobe, superior frontal gyrus - medial orbitofrontal cortex (straight gyrus) |
| lFL,mOFC-SG | left frontal lobe, medial orbitofrontal cortex - straight gyrus |
| lTL | left temporal lobe |
| lTL,sH-LV | left temporal lobe, superior hippocampus - lateral ventricle |
| lTL,sIC | left temporal lobe, superior insular cortex |
| lTL,s | left temporal lobe, superior |
| lTL,m-T2 | left temporal lobe, middle temporal gyrus (T2) |
| lTL,m | left temporal lobe, medial |
| lTL,mH(head) | left temporal lobe, medial hippocampus (head) |

Table A.3 – *Continued from previous page*
| **Label** | **Description** |
| --- | --- |
| lTL,iFG-s | left temporal lobe, inferior fusiform gyrus - superior |

**Table A.4:** Labels describing the anatomical location of bipolar recordings from patient HEC002.

| **Label** | **Description** |
| --- | --- |
| rFL | right frontal lobe |
| rFL,FPO | right frontal lobe, frontal pars opercularis |
| rFL,iFG | right frontal lobe, inferior frontal gyrus |
| rFL,MC(BA6) | right frontal lobe, motor cortex (Brodmann area 6) |
| rTL | right temporal lobe |
| rTL,sAC | right temporal lobe, superior auditory cortex |
| rTL,iIC | right temporal lobe, inferior insular cortex |
| rTL,iaIC | right temporal lobe, inferior anterior insular cortex |
| rTL,H | right temporal lobe, hippocampus |
| rTL,sIC | right temporal lobe, superior insular cortex |
| rTL,IC-FPO | right temporal lobe, insular cortex - frontal pars opercularis |
| rTL,A-B | right temporal lobe, amygdala - brainstem |
| rTL,A-EC | right temporal lobe, amygdala - entorhinal cortex |
| rTL,TP(WM) | right temporal lobe, temporal pole (white matter) |
| rTL,TP(T1-T2) | right temporal lobe, temporal pole (T1-T2) |
| rTL,AC | right temporal lobe, ambient cistern |

Table A.4 – *Continued from previous page*
| **Label** | **Description** |
| --- | --- |
| rTL,H(head)-AC | right temporal lobe, hippocampus (head) - ambient cistern |
| rTL,maH(head) | right temporal lobe, medial anterior hippocampus (head) |
| rTL,H | right temporal lobe, hippocampus |
| rTL,TG(T1-T2)-FG | right temporal lobe, temporal gyrus (T1-T2) - fusiform gyrus |
| rTL,TP-T2 | right temporal lobe, temporal pole - T2 |

**Table A.5:**
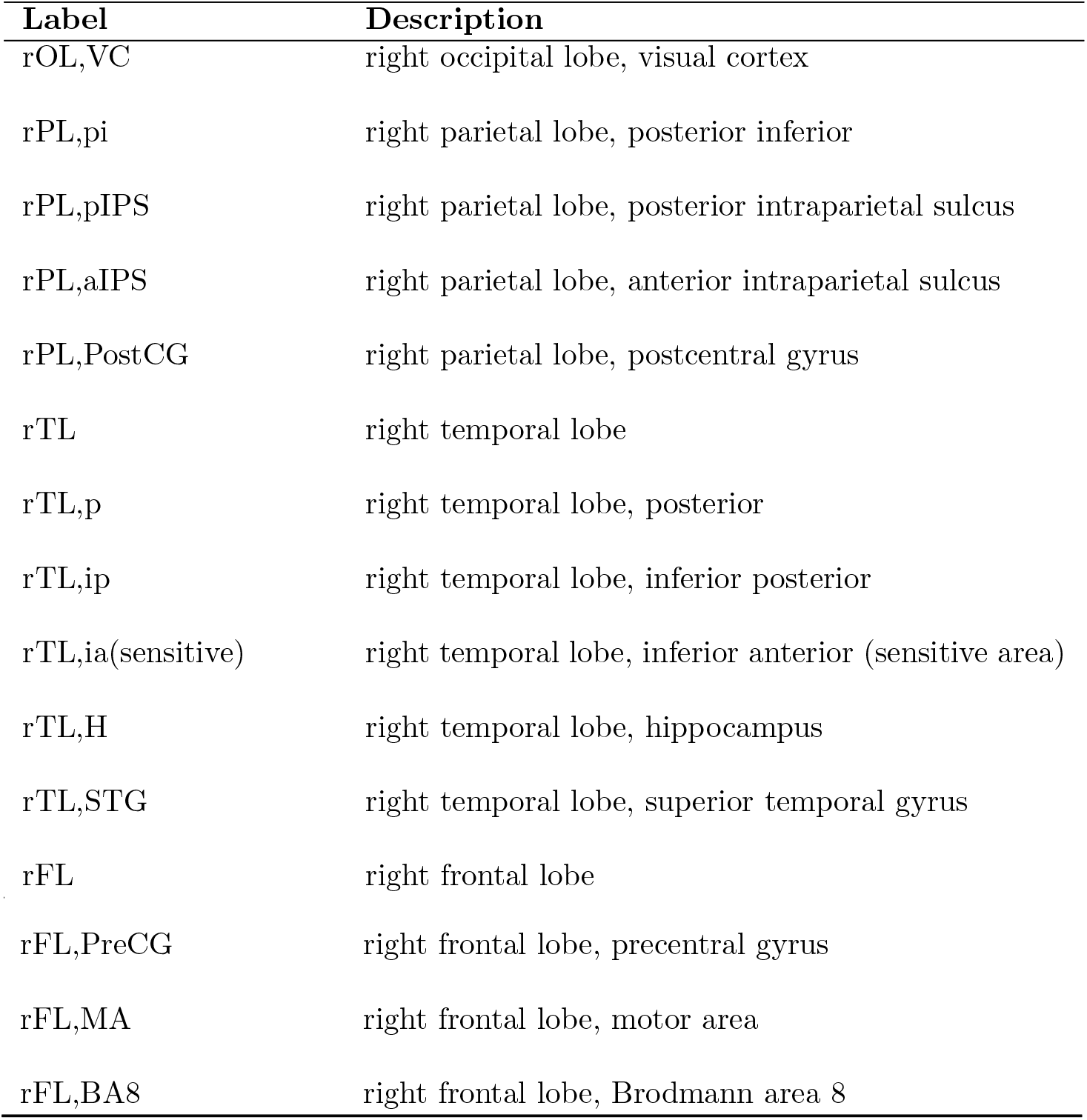
Labels describing the anatomical location of bipolar recordings from patient HRM011.

